# Deep Computational Anatomy via Latent-Aligned Multiview Normalizing Flows

**DOI:** 10.64898/2026.05.05.723039

**Authors:** Nicholas J. Tustison, Brian B. Avants, Philip A. Cook, James C. Gee, James R. Stone

## Abstract

In modeling complex probability distributions, normalizing flows provide exact-likelihood, bijective mappings between empirical data and tractable latent spaces. Building on this foundation, latent-aligned multiview normalizing (LAMNr) flows leverage these salient properties to learn shared latent subspaces across heterogeneous, multimodal datasets while simultaneously topologically unfolding the sampled data manifold into a continuous vector space. Formal latent-alignment constraints are used to model shared structural features separate from view-specific variations, coordinating latent projections into a shared geometric subspace. By applying this transformation in the context of biological imaging, the framework establishes a potential basis for a deep learning interpretation of foundational computational anatomy concepts, such as the population template, latent distances, and geodesic pairwise image interpolation. Additionally, the proposed framework enables closed-form conditional modeling for exact cross-view imputation and other latent space manipulations. Evaluations and illustrations on both imaging-derived phenotypes (IDPs) and multimodal MRI demonstrate the proposed framework and potential applications. To further motivate our work, we provide a robust and comprehensive, 2D and 3D open-source implementation in PyTorch, natively integrated with the ANTsX ecosystem (i.e., ANTsTorch) for efficient training and subsequent data transformation, manipulation, and analysis.

## 1 Introduction

Medical imaging data and their representative latent spaces have become fundamental for insight into biological structure and function. While deep learning has become the current standard for navigating these high-dimensional spaces, certain aspects of contemporary architectures limit exploration, quantitative analysis, and other potential applications through latent space evaluations and manipulations. Generative Adversarial Networks (GANs), for instance, are implicit samplers trained with divergence surrogates rather than likelihoods, which precludes calibration by exact probabilities (Papamakarios et al., 2021). Variational Autoencoders (VAEs) optimize an evidence lower bound rather than the exact log likelihood (Kobyzev et al., 2021). Diffusion and score-based models rely on denoising or score-matching objectives with likelihoods obtained only indirectly (Croitoru et al., 2023). Finally, while autoregressive decoders offer exact likelihoods, they do not yield a one-shot invertible latent representation (Papamakarios et al., 2021). Such limitations extend to multimodal and multiview settings, where heterogeneous or missing data often require cross-view comparisons and coherent anatomical reconstructions.

### 1.1 Normalizing Flows

Normalizing flows model complex data distributions by composing invertible transformations that map input data to their corresponding latents. This bijective design simultaneously yields exact likelihoods via the change-of-variables formula, single-pass inversion, and direct access to latent variables that can be manipulated and decoded without approximation (Kobyzev et al., 2021; Papamakarios et al., 2021). Early developments established the properties and advantages of invertible networks and flow-based density models (Dinh et al., 2017, 2014; Gomez et al., 2017; Jacobsen et al., 2018; Kingma et al., 2016; Papamakarios et al., 2017; Rezende and Mohamed, 2015). Later, Glow architectures introduced data-dependent normalization, invertible 1 × 1(×1) convolutions, and a multiscale structure optimized for imaging (Kingma and Dhariwal, 2018), with subsequent variants improving coupling transforms and stability while preserving exact likelihoods (Chen et al., 2019; Durkan et al., 2019; Grathwohl et al., 2019; Ho et al., 2019). Recent work has demonstrated that flows scale to resolutions and sample qualities comparable to other state-of-the-art generative models (Croitoru et al., 2023; Gu et al., 2025; Zhai et al., 2024). Beyond density estimation, normalizing flows provide a geometric framework for topologically unfolding the complex anatomical manifold sampled by modern medical imaging. In mapping complex imaging data to a symmetric Gaussian base distribution, the flow-induced metric ensures that latent paths approximate geodesics in the original data domain (Arvanitidis et al., 2018; Kobyzev et al., 2021). Recent advancements have further refined these flow trajectories by incorporating Semi-Discrete Optimal Transport (SDOT) during training (Kong et al., 2025). This approach establishes an explicit, optimal alignment between the noise distribution and data points to ensure straighter paths and more effective inference, even in high-dimensional settings.

The bijective formulation of these models also enables the synthesis of biological variation through stochastic sampling, where latent vectors drawn from the Gaussian prior can be mapped back to the high-dimensional image space. While latent diffusion and flow matching achieve high sample quality, they optimize denoising or continuous-transport objectives rather than exact log likelihoods, requiring multi-step sampling or ODE integration (Croitoru et al., 2023; Ho et al., 2020; Lipman et al., 2022). Further distinction between normalizing flows and other generative architectures, such as Generative Adversarial Networks (GANs) and diffusion models, lies in the optimization objective. While the latter often prioritize perceptual-based losses through divergence surrogates or denoising objectives, normalizing flows optimize exact log-likelihoods in latent space. This probabilistic formulation is foundational as it ensures that the latent space functions primarily as a simplified coordinate system with known structure rather than prioritizing aesthetic considerations for image synthesis.

### 1.2 Multiview Learning with LAMNr Flows

Multiview learning operates on two complementary principles: 1) each distinct acquisition or feature space (“view”) contributes unique, view-specific information and 2) shared information across views can be transformed via projections (often to lower-dimensional space) to improve calibration and cross-cohort comparability. Traditionally, these shared projections have been estimated using classical correlation-based methods, such as Canonical Correlation Analysis (CCA) (Hardoon et al., 2004; Hotelling, 1936). More recently, kernel-based measures, like the Hilbert–Schmidt Independence Criterion (HSIC) (Gretton et al., 2005), and learned alignment objectives, including Barlow Twins (Zbontar et al., 2021), VICReg (Bardes et al., 2022), and InfoNCE (Oord et al., 2018), have expanded these capabilities to accommodate more complex patterns.

One example, similarity-driven multilinear reconstruction (SiMLR), captures this joint variation in a linear, low-rank setting by projecting multiview data into a shared subspace under subject-level similarity constraints (Avants et al., 2021). While SiMLR isolates this shared representation, it treats the remaining view-specific variation as an unstructured residual rather than explicitly modeling a private component. This separation supports robust cross-view harmonization and prediction by isolating stable population effects from noise (Stone et al., 2024, 2020). While deep learning approaches have explored cross-modal translation and disentanglement using convolutional neural networks, VAEs, or diffusion models (Chartsias et al., 2019; Havaei et al., 2016; Yuan et al., 2024), they often lack the unique combination of exact likelihoods and one-shot invertible mappings. Recent work has also explored normalizing flows for unsupervised MRI harmonization, but utilize the flow purely as a test-time density estimator to iteratively adapt an auxiliary translation network to an unknown target domain (Beizaee et al., 2025).

In contrast, LAMNr flows analogize the SiMLR framework into a deep, likelihood-based architecture that topologically unfolds the potentially complex manifold into a continuous vector space. Instead of an explicit linear factorization in the observation domain, LAMNr flows map each view into a shared latent space, ensuring exact log-likelihoods and bijective mappings. By utilizing latent-alignment objectives (e.g., VICReg, InfoNCE) to identify shared coordinates, the framework recovers the interpretability of a shared/private decomposition within a nonlinear, invertible space. Crucially, by modeling the joint latents with a Gaussian distribution, LAMNr flows enable closed-form conditional reconstructions. This allows the shared subspace to function as a geometrically-informed coordinate system. Recent efforts have explored multiview-enriched normalizing flows for complex density estimation (Kruse and Rosenhahn, 2025) using standard RealNVP architectures. LAMNr flows specifically extend this logic by integrating a latent shared/private decomposition that enables geometrically-informed coordinate systems for multimodal alignment while also including image-specific Glow architectures.

Additionally, the development of LAMNr flows provides a practical strategy in ensuring topological integrity within neural density estimators. Historically, models like Deep Diffeomorphic Normalizing Flows (DDNF) (Salman et al., 2018) enforced smoothness by integrating time-varying velocity fields via Ordinary Differential Equations (ODEs). While this continuous formulation guarantees a diffeomorphic mapping, the computational cost of ODE integration is often prohibitive for largescale applications (e.g., medical imaging). To address this, LAMNr flows use the discrete, efficient architectures of RealNVP (Dinh et al., 2017) and Glow (Kingma and Dhariwal, 2018). By aligning disparate modalities and views into a shared latent representation, the LAMNr flows model is steered towards prioritizing robust, underlying anatomical structures over idiosyncratic signal. Additionally, this latent-alignment employs specific numerical safeguards, such as bounding the scale parameters within the affine coupling layers, to mitigate gradient blow-ups during training. Furthermore, the inclusion of training jitter serves as an additional regularizer (i.e., “dequantization” (Ho et al., 2019)). In the imaging context (i.e., Glow-based models), by introducing stochastic intensity-based and shape-based perturbations during the learning phase, the model is discouraged from overfitting to local voxel intensities. Together, these constraints force convergence on more generalized anatomical representations, facilitating the stabilization of the Jacobian determinant and ensuring that the discrete transitions of the Glow architecture maintain the smooth, diffeomorphic properties required. Analogous network architectural features are also leveraged for IDP-based scenarios using RealNVP.

### 1.3 Computational Anatomy and Normalizing Flows

Computational anatomy (CA) is a comprehensive mathematical discipline that formalizes the study of biological shape and its variability through the action of diffeomorphic transformation groups on anatomical manifolds (Grenander and Miller, 1998; Miller et al., 2002). As one important example, within this probabilistic and geometric framework, population structure is typically represented by a population template, i.e., a reference space formally established as the Fréchet mean that minimizes the sum of squared geodesic distances across a cohort (Avants et al., 2010). Traditionally, the intrinsic curvature of image spaces causes a divergence between the Fréchet mean, the Karcher mean, and the statistical mode (Fletcher et al., 2009).

Normalizing flows offer an alternative, deep learning perspective by topologically unfolding these nonlinear manifolds into a symmetric, centered diagonal Gaussian base distribution (Figure 1). Within this framework, one principled approach to template construction is the inverse mapping of the latent origin, 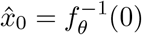, which leverages the property that the Gaussian mean, mode, and median coincide precisely at the origin (cf. Figure 1). While registration-based templates (e.g., via Symmetric Normalization (Avants et al., 2010) or Large Deformation Diffeomorphic Metric Mapping (Miller et al., 2002)) typically preserve high-frequency details through iterative normalization, spatial averaging, and sharpening, the generative latent origin-based template exhibits a visually smoother appearance. This smoothness is a direct consequence of high-dimensional probabilistic modeling. As the exact mode of the latent distribution, the origin averages out idiosyncratic, high-frequency anatomical variations, such as individual cortical folding patterns, that do not strictly persist across the cohort.

**Figure 1:**
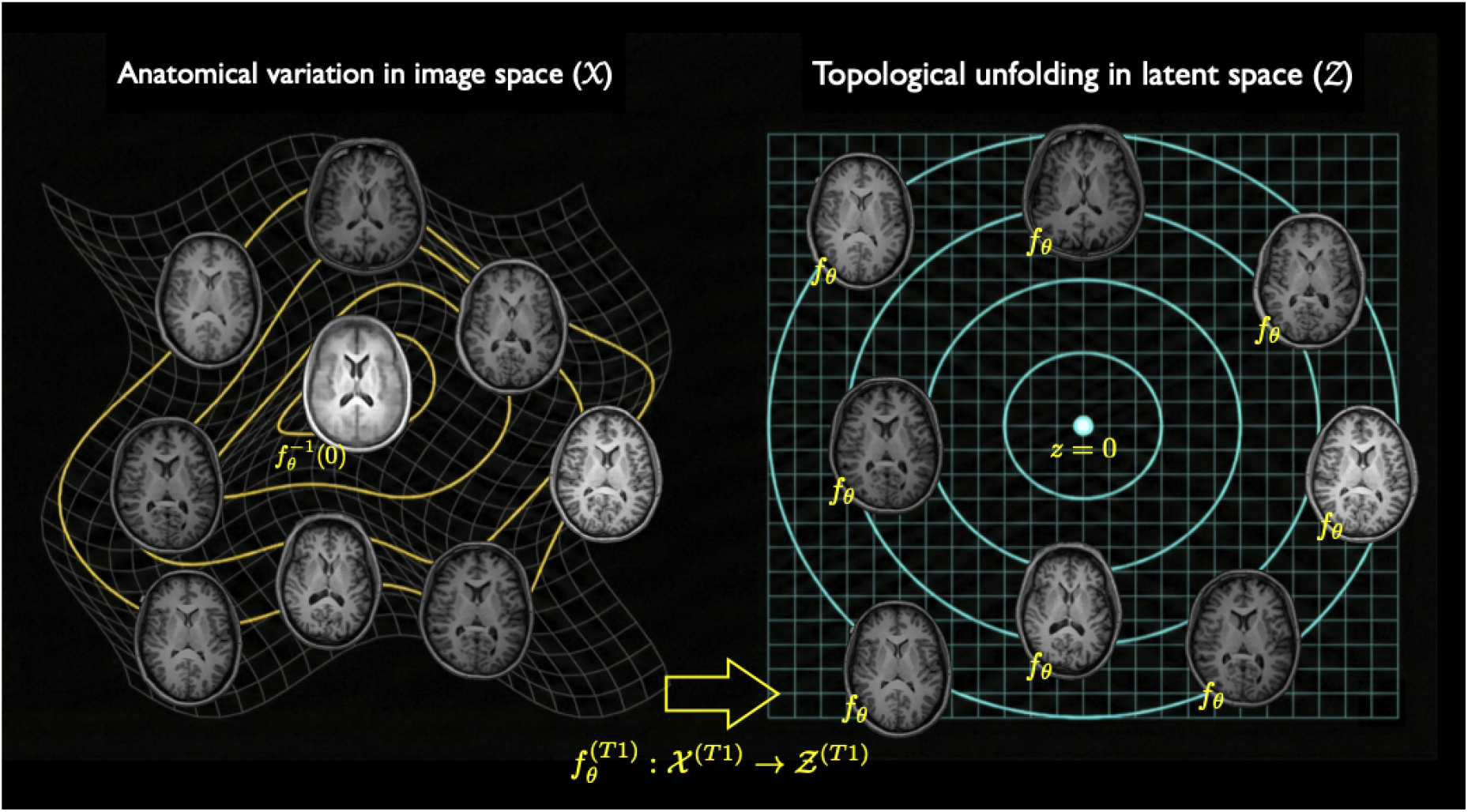
Bijective mapping between image and latent spaces established by a normalizing flow for a single modality (e.g., T1-weighted MRI). The architecture projects complex anatomical data from the observation space 𝒳 (left) to a tractable, symmetric Gaussian base distribution in the latent space Ƶ ~ 𝒩 (0, 1) (right). This single-pass encoding and decoding mechanism provides the exact-likelihood foundation upon which the LAMNr flows framework is built.

**Figure 2:**
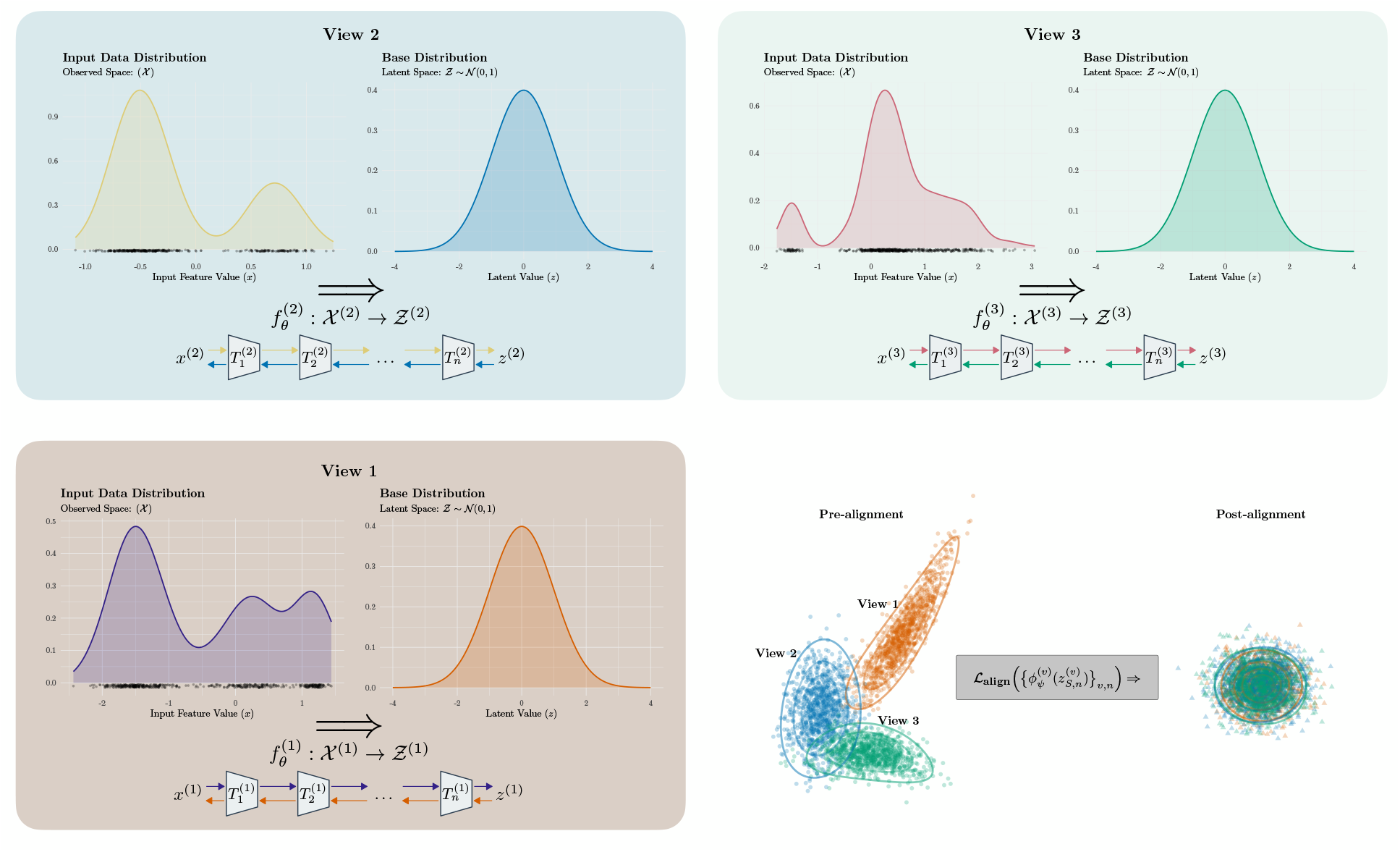
LAMNr flows architecture and latent alignment. The model processes three data views, illustrated by the panels View 1, View 2, and View 3. For each view, the data distribution in the observation space 𝒳 can be mapped to a simplified base distribution in the latent space Ƶ ~ 𝒩 (0, 1). This mapping is performed by the individual normalizing flows sequential bijections (*T*_1_, *T*_2_, …, *T*_*n*_). Joint alignment optimization is performed on latent distributions to drive convergence towards a normalized shared space through the application of the alignment loss function 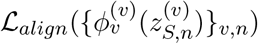.

Furthermore, regarding the geometric interpretation of this template, in high dimensions the concentration of measure phenomenon dictates that probability mass concentrates within a thin spherical shell rather than at the origin (Blum et al., 2020; Vershynin, 2018; White, 2016). While classical geometric morphometry often utilizes the Fréchet mean to compute a statistically valid average on a Riemannian manifold (Pennec, 2006), applying such spherical mapping in a spatial normalizing flow to force the template onto the high-probability “typical set” destroys the anatomical signal. Because LAMNr flows preserve spatial dimensions, projecting the vector norm to this spherical shell normalizes the spatial contrast energy, resulting in severe high-frequency noise. Consequently, the latent origin *z* = 0 is not a statistically typical anatomical instance, but rather a barycentric geometric anchor representing the central axis of symmetry for the learned bijection.

Beyond template construction, this continuous latent framework provides direct analogues to the fundamental metric operations of traditional computational anatomy. For example, in classic diffeomorphic frameworks, the transformation between a source and target anatomy is governed by integrating a time-varying velocity field over a continuous time domain *t* ∈ [0, 1]. The length of this optimal, continuous deformation path establishes the exact geodesic distance between the two biological structures (Beg et al., 2005; Miller et al., 2002). In the LAMNr flows framework, this computationally intensive temporal integration can be substituted with an algebraic interpolation within the latent space. Traversing the latent manifold between two encoded images, *z*_0_ and *z*_1_, using a scalar interpolation parameter *α* ∈ [0, 1] generates a continuous trajectory of decoded images that closely approximates this diffeomorphic flow. Consequently, the distances computed directly in the latent space, when properly evaluated via distribution-preserving spherical metrics rather than Euclidean norms, serve as highly efficient surrogates for the complex, deformation-based geodesic distances of traditional computational anatomy. These conceptualizations, along with other illustrative results, are discussed and provided below in the context of our proposed LAMNr flows framework.

### 1.4 Contributions

We introduce LAMNr flows, a general framework for deep computational anatomy that learns shared and private latent structures across multiple views while preserving exact likelihoods and invertibility. Within the LAMNr flows framework, each view is equipped with a dedicated flow that maps observations to a structured latent space. Key contributions of this work include:

- **Unified Multiview Modeling**. We provide a shared coordinate system for heterogeneous data types, including 2D/3D images and tabular imaging-derived phenotypes (IDPs).
- **Latent Alignment and Linearization**. Using subject-matched batches, we identify shared anatomical features via a library of alignment losses, such as VICReg and InfoNCE.
- **Demonstration of Deep Computational Anatomy**. We demonstrate essential capabilities including closed-form cross-modal data imputation via the Woodbury identity and high-fidelity image interpolation using spherical linear interpolation (Slerp) to prevent variance collapse.
- **Open-source Implementation**. We provide a comprehensive, open-source 2D and 3D PyTorch implementation based on the normflows library. This framework is natively integrated with the ANTsX ecosystem via ANTsTorch for robust data handling and auxiliary functionality.

## 2 Methods

### 2.1 Normalizing Flows and the LAMNr Flows Multiview Formulation

Given subject *n* with measurements 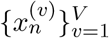 across one or more views *v* = 1, …, *V*, for a single view *v*, a normalizing flow with parameters *θ*^(*v*)^ is an invertible mapping

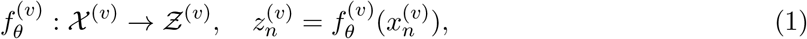

that transforms observed data 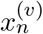 to a latent space Ƶ ^(*v*)^ with a chosen base density, typically *p*_*Z*_(*z*) = 𝒩 (0, 1). The induced density on 𝒳 ^(*v*)^ follows from the change-of-variables formula:

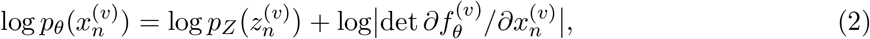

which can be evaluated exactly for the RealNVP (Dinh et al., 2017) and Glow (Kingma and Dhariwal, 2018) architectures. The maximum-likelihood estimation chooses *θ*^(*v*)^ to maximize the sum of log-likelihoods over subjects, or equivalently to minimize the average negative log-likelihood (NLL) (Kobyzev et al., 2021).

For multiple views in the LAMNr flows framework, we instantiate one flow per view and train them jointly on subject-matched minibatches. If we consider only the flow likelihoods, the pure maximum-likelihood objective is

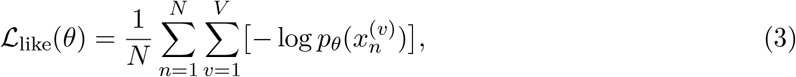

where 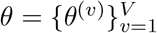 represents all view-specific parameters. This term ensures that each per-view flow is an exact-likelihood model of its corresponding data distribution. For each view and subject we write

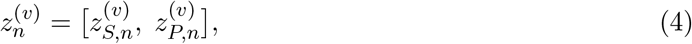

splitting the latent into a block 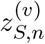 that is intended to carry shared information across views and a block 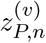 that is view-specific. The indices that define this split can be chosen a priori or via the CCA/HSIC-based screening procedure (described below).

We attach a small projector network 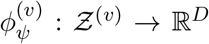 to each view, utilizing a multi-layer perceptron (MLP) architecture with a hidden layer width *H* and a matched output dimensionality *D* across all views. Default values are *H* = 512 and *D* = 256. Attaching this projector decouples the flow’s internal latent dimensionality and arbitrary coordinate system from the alignment space. This allows each view to learn a light reparameterization of alignment of rotations and scales frequently introduced by invertible mixing layers such as the 1 × 1(×1) convolutions in Glow, while harmonizing dimensions across disparate views. The role of this *D* subspace can vary strategically depending on the data type. For tabular IDPs (i.e., RealNVP) this configuration represents a high-capacity expansion of the lower-dimensional latent space. This expansion provides the alignment constraints with sufficient degrees of freedom to operate without inducing information loss or architectural bottlenecks. For image data (i.e., Glow), the projection acts as an intentional dimensionality reduction and selective filter. By compressing the massive raw latent space into the lower space *D*, we filter out view-specific, high-frequency noise, ensuring the alignment objective captures shared, global morphometric trends rather than idiosyncratic imaging artifacts. By restricting alignment to this matched subspace, we stabilize the specified alignment constraint and avoid the instability of forcing private, view-specific anatomical directions to align. We summarize the main options in Table 1. In all cases, the alignment term acts only on these shared coordinates, leaving private coordinates free to capture independent variation.^1^

**Table 1:** Summary of latent alignment objectives used in LAMNr Flows.

| Method | Objective | Hyperparameters | Qualitative behavior / usage |
| --- | --- | --- | --- |
| **Pearson (multi)** | Maximize mean pairwise correlation: $\mathcal{L} = \text{corr}(\tilde{Z}^{(i)}, \tilde{Z}^{(j)})$ | Projector dimension; feature normalization (pre-BN / L2) | Emphasizes linear shared structure; simple, fast, and stable at small batch sizes; purely second-order, can over-smooth fine texture if heavily weighted at high levels. |
| **Barlow Twins (multi)** | Cross-correlation to identity: $\mathcal{L} = \sum_i (1 - C_{ii})^2 + \lambda \sum_{i \neq j} C_{ij}^2$ | $\lambda$ (off-diagonal weight); covariance shrinkage; projector dimension | Encourages invariance and decorrelation; no negatives; good default when moderate batch sizes are available; shrinkage improves stability of $C$. |
| **VICReg (multi)** | $\mathcal{L} = \alpha \text{Inv} + \beta \text{Var} + \gamma \text{Cov}$ | $\alpha, \beta, \gamma$; variance margin; projector dimension | Promotes invariance while preserving variance and discouraging collapse; works with small-to-moderate batches; more tuning knobs (trade-off between invariance, variance, and decorrelation). |
| **InfoNCE (multi)** | Contrastive: $\mathcal{L} = -\log \frac{\exp(s/\tau)}{\sum \exp(s'/\tau)}$ | Temperature $\tau$; projector dimension; (optional) augmentation strength | Strong discriminative alignment when large batches and many in-batch negatives are available; higher compute cost; sensitive to batch size and choice of $\tau$. |
| **HSIC (biased)** | Maximize kernel dependence: $\mathcal{L} = \text{HSIC}(X, Y)$ (e.g., RBF kernel) | Kernel type; bandwidth $\sigma$ (median heuristic); regularization | Captures non-linear shared structure beyond correlations; useful when complex cross-view relations are expected; $\mathcal{O}(B^2)$ cost in batch size and sensitive to bandwidth choice. |

**Legend:** $\tilde{Z}^{(i)}$ and $\tilde{Z}^{(j)}$ are batch-normalized projector outputs for two views; $C$ is their cross-correlation matrix with entries $C_{ij}$ ; Inv, Var, and Cov denote the invariance, variance, and covariance terms in VICReg; $s$ and $s'$ are similarity scores between paired and non-paired samples; $\tau$ is the temperature; $X$ and $Y$ are projected features from two views used to compute $\text{HSIC}(X, Y)$ ; and $\sigma$ is the kernel bandwidth.

To prevent over-constraining the flows and blurring view-specific information, we perform a short screening pass after an initial “warm-up’ ‘ phase. For CCA-based screening, we construct whitened feature matrices for two views and perform an SVD of the cross-covariance 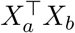, retaining the top *r* canonical directions per view. Averaging across all view pairs yields per-view projectors *P* ^(*v*)^ ∈ ℝ^*D*×*r*^ defining the shared subspace. For HSIC-based screening, we first prefilter coordinates using Pearson correlation, then rank remaining dimensions by an unbiased HSIC estimate with RBF kernels averaged over other views, and select the top *r* dimensions per view. Alignment losses are applied only to these projected or masked coordinates, so that dependence is enforced where cross-view signal is strongest and private dimensions remain free to capture view-specific variation. Screening can be performed once after warm-up or periodically refreshed during training. For a subject-matched minibatch of size *N*, the full training objective becomes

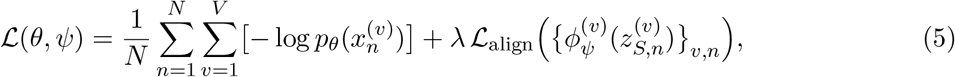

where ℒ_align_ is one of the available options: Barlow Twins, VICReg, InfoNCE, Pearson correlation, or HSIC. *λ* controls the strength of alignment.^2^

### 2.2 View-specific Flow Architectures

#### 2.2.1 Tabular/IDP Views via RealNVP

For imaging-derived phenotypes (IDPs) and other tabular data, we use single-scale flows based on RealNVP and masked autoregressive flows (MAF) with affine couplings and masked multilayer perceptrons. Continuous variables are preprocessed per view using dataset-owned normalization and imputation: columns are coerced to numeric, NaNs are imputed as the column mean, and features are standardized to zero mean or rescaled to [0, 1] depending on a user-selectable normalization mode. Very low-variance columns are stabilized by floor-clamping the standard deviation. Positively skewed, non-negative variables can optionally be log- or log 1*p*-transformed before normalization to reduce skewness.

We use two base distributions: a diagonal Gaussian and a Gaussian–PCA base that performs an additional linear whitening of the flow latents. In the latter case, the flow acts as a learnable multiview “whitener” that maps each tabular view to a standardized latent *ε* with approximately independent components. Both the raw flow latents *z*^(*v*)^ and the whitened coordinates *ε*^(*v*)^ can be exported for downstream Gaussian modeling and diagnostics. This Gaussian–PCA base distribution is particularly useful when tabular views have different numbers of features. The per-view PCA yields an orthonormal, variance-ordered latent in which we can select a common rank *r* for alignment, producing matched-dimension standardized coordinates *ε*^(*v*)^ ∈ ℝ^*r*^ without altering the exact invertibility of the flow as truncation is used only for the alignment head.

#### 2.2.2 Image Views via Glow-based Multiscale Flows

For image views we adopt Glow-style discrete normalizing flows with *L* levels and *K* coupling steps per level and a diagonal Gaussian base distribution. Each coupling step comprises: (i) ActNorm layers with data-dependent initialization, (ii) invertible 1 × 1(×1) convolutions parameterized with LU factorization for efficient log-determinant computation, and (iii) affine coupling layers whose scale and shift fields are predicted by shallow convolutional subnetworks with a configurable number of hidden channels. Squeeze and split operations provide a multiscale representation in which shallower levels capture coarse structure while deeper levels model fine texture. Our implementation follows the standard Glow construction, instantiated via a model factory in ANTsTorch, with configurable image size (both 2D and 3D), number of levels *L*, steps per level *K*, and number of hidden channels.^3^

For image views we use a channel-wise diagonal Gaussian (“Glow base”) with one mean and one log-scale per channel, broadcast across spatial locations. Let 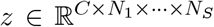 with *S* ∈ {2, 3} spatial dimensions and 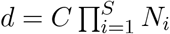. The log density is

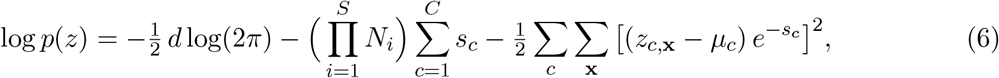

where *µ*_*c*_ and *s*_*c*_ are per-channel parameters broadcast over all spatial indices **x**. Compared to a conventional per-voxel diagonal Gaussian, tying parameters within each channel reduces degrees of freedom, matches Glow’s multiscale semantics, and avoids per-voxel scale collapse.

### 2.3 High-dimensional Geometry and Latent Space Navigation

In high-dimensional standard normal latent spaces, such as those optimized by LAMNr flows, the geometric properties of the data distribution become counterintuitive due to the concentration of measure phenomenon (Blum et al., 2020; Vershynin, 2018; White, 2016). In other words, as dimensionality increases, probability mass moves away from concentration at the origin. Instead, the volume of the space grows exponentially with distance from the center, causing the vast majority of the mass to concentrate within a narrow spherical shell of radius 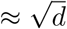 (i.e., the “soap bubble” effect^4^), a consequence of the Gaussian Annulus Theorem (Blum et al., 2020), often referred to as the typical set (Blum et al., 2020; Vershynin, 2018). Consequently, the latent origin *z* = 0 is a highly atypical point containing near-zero probability mass. The inverse mapping *f*^−1^(0) must therefore be understood strictly as a barycentric geometric anchor representing a central axis of symmetry for the learned bijection, rather than a statistically representative anatomical mode.

Furthermore, as the normalizing flow unfolds the global topology of the anatomical data, the resulting latent space is not a flat Euclidean manifold such that Euclidean operations in the latent space do not translate to valid anatomical transformations in the image space. Rather, latent-based distances are governed by a stochastic Riemannian metric induced by the generator’s Jacobian, defined as 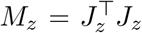 (Arvanitidis et al., 2018). Because the network non-linearly expands and compresses the data space to maximize likelihood, Euclidean straight lines in the latent space do not correspond to the shortest paths (geodesics) on the underlying image manifold. This geometric distortion has immediate, tangible consequences for cohort alignment and interpolation. Linearly interpolating between two latent points located on the typical set creates a trajectory that moves inward toward the latent origin. In high dimensions, this effect forces the interpolation path through unpopulated latent regions of extremely low probability, causing a distribution mismatch (Agustsson et al., 2018). The resulting generated images exhibit blurriness, structural artifacts, and anatomical inconsistencies.

To navigate this geometry appropriately, we replace standard linear interpolation with spherical linear interpolation when traversing the latent space between two generated samples *z*_1_ and *z*_2_ (Agustsson et al., 2018; White, 2016). For an interpolation parameter *t* ∈ [0, 1] and the angle 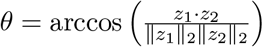 between the vectors, the Slerp (i.e., spherical linear interpolation) trajectory is defined as:

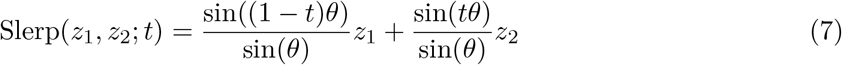

This formulation ensures that the interpolation path strictly follows the high-probability manifold, preserving structural integrity. Similarly, we adapted our distance metrics based on the evaluation context. When assessing the semantic similarity between two individual images within the latent space, we default to the geodesic (angular) distance rather than the Euclidean distance. The geodesic distance effectively isolates the directional components of the vectors:

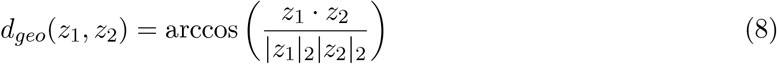

This metric captures core semantic features while discarding magnitude variations that primarily represent high-dimensional statistical noise. However, for measuring a subject’s deviation from the normative population, we utilize the Mahalanobis distance relative to the Gaussian mean (*µ* = 0):

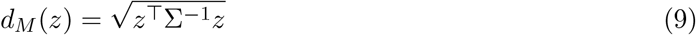

where Σ represents the covariance matrix of the reference cohort.

### 2.4 Conditional Gaussian Modeling Over Latents

Normalizing flows yield an explicit bijection between data and latent spaces with a simple base density (e.g., Gaussian). Once the per-view flows have been trained, every multiview observation 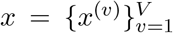 can be mapped to a collection of latents 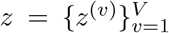 with an exactly known Jacobian. Any joint distribution placed on these latents induces a valid joint distribution on the original data via the change-of-variables formula. In other words, specifying a model *p*_*Z*_(*z*) in latent space is equivalent to specifying a generative model *p*_*X*_(*x*) in data space, but with the advantage that inference and conditioning can be carried out where the geometry is simpler.

LAMNr flows exploit this by choosing a multivariate Gaussian model on the concatenated latents. This choice is deliberately simple as the flows absorb the complex, non-Gaussian aspects of each view into the invertible mappings *f*^(*v*)^, so that the residual cross-view structure can be captured by a Gaussian dependence model in *z*. Under this construction, the joint density factorizes as

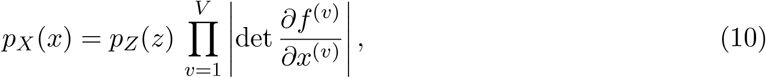

with *z* = *f* (*x*). Because *p*_*Z*_(*z*) is Gaussian, all conditionals *p*_*Z*_(*z*_*U*_ | *z*_*O*_) are available in closed form, and exact conditional inference in data space reduces to three steps: 1) encode observed views to latents, 2) apply Gaussian conditioning in *z*, and 3) decode the resulting latents back through the inverse flows. This yields closed-form posteriors, imputations, and counterfactuals that are fully consistent with the learned flow model.

After training the per-view flows and projector alignment, we freeze the flow parameters and collect latents for all subjects. For image views, we retain a multiscale representation 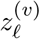 at each level *ℓ* ∈ {1, …, *L*} whereas for tabular views, we have a single level. Concatenating across views and levels yields a joint latent vector

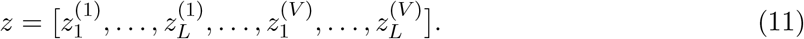

We model this joint latent as Gaussian, *z* ~ 𝒩 (*µ*, Σ). However, directly computing and operating on the full covariance matrix Σ poses a severe computational bottleneck due to the “curse of dimensionality.” While feasible for 2D images, a small 3D medical volume (e.g., 64 × 64 × 64) yields a latent dimension *D* ≈ 2.6 × 10^5^. Storing the dense *D* × *D* covariance matrix as 64-bit double precision requires over 500 GB of memory, making direct Cholesky inversion computationally intractable.

To resolve this in high-dimensional settings, we employ a low-rank-plus-diagonal parameterization via Singular Value Decomposition (SVD). We approximate the covariance as Σ ≈ *U* Λ*U*^*T*^ + *σ*^2^*I*, where *U* contains the top *r* eigenvectors (*r* ≪ *D*), Λ is the diagonal matrix of the top *r* eigenvalues, and *σ*^2^*I* captures the isotropic residual variance. Given an observed subset of coordinates *O* and an unobserved subset *U*, the exact posterior *p*(*z*_*U*_ | *z*_*O*_) is Gaussian. The conditional mean and covariance are theoretically given by:

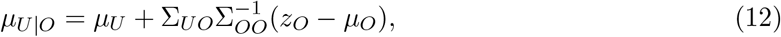

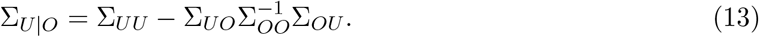

To evaluate these equations without instantiating the massive Σ_*OO*_ matrix, we utilize the Woodbury matrix identity (Henderson and Searle, 1981). Specifically, we apply the Push-Through identity to perform the inversion strictly within the lower-dimensional subspace *r* (Rasmussen and Williams, 2006). By reformulating the system to solve for a strictly positive-definite *r* × *r* matrix 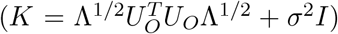, we bypass problematic numerical cancellations (Higham, 2002) and significantly reduce the memory requirements.

Critically, the fidelity of these conditional results is intrinsically linked to the inter-view latent alignment established during the initial training phase. In the joint Gaussian model, the cross-covariance terms Σ_*UO*_ encapsulate the statistical coupling between the observed and unobserved manifolds. Effective alignment ensures that subject-specific anatomical features are mapped to shared latent coordinates, enabling the conditional mean to recover missing modalities with high individual specificity. In the absence of such alignment (i.e., when Σ_*UO*_ ≈ 0), the conditional mean *µ*_*U*|*O*_ collapses to the population prior *µ*_*U*_. In this limit, the imputed output reverts to a generic population template, losing the subject-matched morphological details. Samples from this conditional Gaussian propagate uncertainty, while the posterior mean provides a calibrated point estimate. Applying the inverse flows to these posterior latents yields imputations, harmonized representations, and latent edits in the original data space, with exact likelihoods available for all configurations.

### 2.5 Implementation and Training Details

Our implementation builds on the `normflows` PyTorch package for normalizing flows (Stimper et al., 2023), which we have extensively modfiied for improvements in training normalizing flows and to accommodate the LAMNr flows framework. At the architectural level, we reconfigured the layer ordering to match Glow-style multiscale flows (ActNorm → invertible 1×1(×1) convolution → affine coupling). We also implemented 3D variants of the core components (squeeze / unsqueeze, split / merge, invertible 1×1×1 convolutions, and 3D coupling networks) to support volumetric image data. These models are exposed through ANTsTorch as configurable factories for both image and tabular views: Glow-style flows for 2D/3D images and RealNVP-style flows for IDPs and other tabular blocks. The ANTsTorch interface handles dataset-level normalization and imputation, Gaussian and Gaussian–PCA whiteners with optional application at train and test time.

Several empirical and survey works have noted that, compared to VAEs and diffusion models, deep normalizing flows can be numerically sensitive on high-dimensional data, with stability depending strongly on architectural choices, scale parameterization, and Jacobian conditioning (Behrmann et al., 2019; Croitoru et al., 2023; Durkan et al., 2019; Kobyzev et al., 2021; Papamakarios et al., 2021). In light of this, we introduced several additional stability-oriented modifications in both our ANTsTorch builders and our `normflows` fork. These include bounded coupling scales (via configurable `scale_map` and `scale_cap` parametrizations), constrained base log-scales for Glow-style bases, optional ActNorm inside coupling subnetworks, and gradient-norm clipping. We also refactored computation of the Gaussian log likelihood for the base Σ = *W*^⊤^*W* + *σ*^2^*I* using a Cholesky factorization with a small adaptive jitter, evaluate log |Σ| as 2 Σ log diag(*L*), and form the quadratic term via triangular solves rather than explicit matrix inversion. This avoids determinant and matrix-inverse calls that are unstable in high dimensions and yields fewer NaNs during training. We initialize *W* at small scale so that Σ is well conditioned at start, and we optimize log *σ* to keep *σ* strictly positive. These choices follow standard numerical recommendations for stable positive-definite computations (Higham, 2002) and pair naturally with shrinkage used elsewhere in our conditional-Gaussian step (Ledoit and Wolf, 2004; Schäfer and Strimmer, 2005). Collectively, these changes reduce log-det explosions and latent outliers in deep multiscale flows while preserving exact likelihoods and invertibility.

Training and validation splits are defined at the subject level, and each minibatch contains aligned multiview data from matched subjects. Image data augmentation is performed on-the-fly using the ANTsTorch-based `ImageDataset` with affine and diffeomorphic deformations, small intensity perturbations (histogram warping and bias field simulation (Tustison et al., 2021a)), and additive Gaussian noise treated as dequantization (Ho et al., 2019) rather than biological variability. We control the overall augmentation strength by a scalar schedule *α*(*t*) ∈ [0, 1] as a function of normalized training time *t*, and support linear, cosine, and exponential decay. For example, a linear schedule reduces augmentation proportionally to *t*, a cosine schedule keeps stronger perturbations early and then decays smoothly, and an exponential schedule reduces aggressive warps and noise most rapidly at the beginning of training. This allows us to start with heavier augmentations to regularize the flows and discourage overfitting to discrete templates, then gradually emphasize fidelity to the true data distribution as training progresses. This design preserves anatomical variability while preventing overfitting to discrete, noise-free templates that would otherwise cause flows to collapse onto certain background modes.

#### 2.5.1 Tabular-specific Implementation Details

For tabular flows, we apply a small additive “jitter” noise to the features, treated as dequantization (Ho et al., 2019) rather than biological variation. The amplitude of this jitter is managed by a scalar schedule *α*(*t*), supporting linear, cosine, or exponential decay, to regularize the flows and prevent overfitting to discrete patterns or exact repeated rows in large cohorts. In addition, certain views can undergo a per-feature marginal transform prior to normalization, such as an elementwise asinh(*x*) for heavy-tailed continuous variables or rank-based Gaussianization that maps the empirical CDF of each feature to a standard normal. These monotone transforms preserve rank information while making marginals more Gaussian and reducing extreme tails.

The underlying training architecture, as implemented in `train_lamnr_flows_tabular.py`, facilitates a highly modular approach to latent alignment and density estimation. Users can choose between a standard DiagGaussian base or a GaussianPCA distribution, which acts as a learnable, geometrically-informed coordinate system to perform linear whitening of the flow latents. This is particularly useful for multiview alignment, as it allows for the selection of a common rank *r* across disparate views. To further refine the alignment process, the framework includes an optional pre-training screening pass based on Canonical Correlation Analysis (CCA) or HSIC. This screening evaluates cross-view dependence to determine if alignment constraints (e.g., VICReg, HSIC, or InfoNCE) should be active for specific view pairs, thereby preventing the model from over-constraining the flows and blurring view-specific information. Numerical stability across these operations is ensured through bounded coupling scales and ActNorm layers, providing a robust foundation for identifying shared anatomical signals.

#### 2.5.2 Glow-specific Implementation Details

The training interface for imaging views, implemented in `train_lamnr_glow_2d.py` and `train_lamnr_glow_3d.py`, utilizes a GlowStepWrapper to parallelize exact log-likelihood and latent prior computations. Models are initialized with data-dependent ActNorm, followed by a one-time warm-up pass with real images to stabilize statistics. Optimization is performed using Adamax with mixed precision (GradScaler) and a learning rate schedule that includes both a warm-up phase and a plateau-based reducer. To improve generative stability and sample quality, we also maintain an Exponential Moving Average (EMA) of model parameters.

To accommodate training scenarios under constrained VRAM, particularly for 3D volumes, we enable gradient accumulation (microbatching). With an accumulation factor *A* and microbatch size *B*_*µ*_, the effective batch size is *B*_eff_ = *A · B*_*µ*_. We compute per-sample losses on each microbatch, accumulate their gradients, and perform a single optimizer step every *A* microbatches. Likelihood terms are summed in natural units and normalized by the total number of samples across the *A* microbatches. Alignment losses (specifically VICReg and InfoNCE) are normalized by the number of accumulation steps to ensure gradient consistency with the non-accumulated baseline. While this preserves the scale of the objective, the alignment constraints are computed within each micro-batch rather than across the full effective batch. Consequently, the contrastive “negative pool” and covariance statistics are derived from local batch statistics, providing a robust but computationally feasible regularization of the manifold. Under mixed precision, gradients are accumulated in scaled form and unscaled once before a single global-norm clip and optimizer step. EMA updates and learning-rate schedulers advance once per effective batch, while ActNorm layers utilize fixed statistics after the initial warm-up phase to ensure that accumulation does not alter normalization behavior.

To ensure stable density estimation and prevent degenerate likelihoods due to data quantization, we also employ uniform dequantization (jittering) during training, following the variational framework established in Flow++ (Ho et al., 2019). We apply lightweight, label-free augmentations during maximum-likelihood training to improve robustness without changing the model’s exact likelihood computation (augmentations act on inputs only) (Tustison et al., 2025, 2024). For image views, we use geometric transforms (linear and non-linear transformations) shared across all views of a subject to preserve alignment targets, and per-view intensity-based transforms (noise, simulated bias-field, histogram warping). Similar to the tabular case, the amplitude is controlled by a scalar schedule *α*(*t*) (linear, cosine, or exponential in training time) (cf Figure 3).

**Figure 3:**
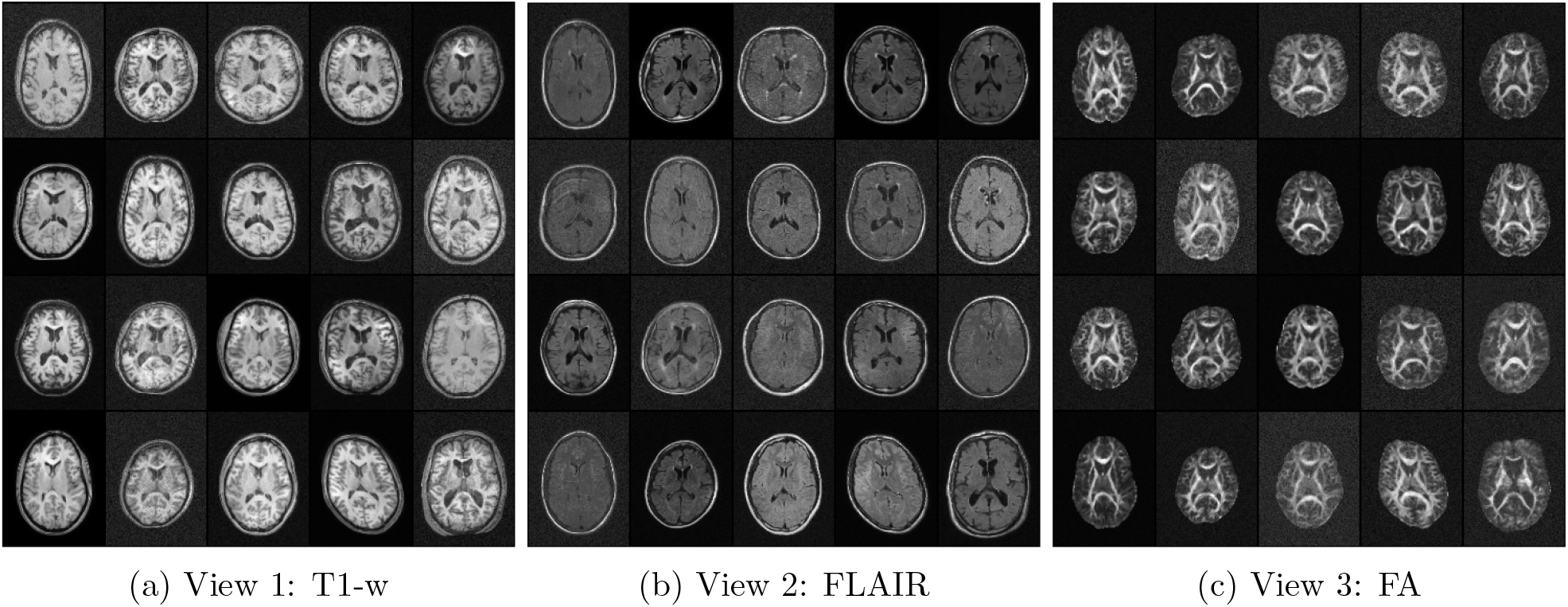
Illustration of the coordinated data augmentation across modalities. Shown are augmented mid-axial slices from the DLBS dataset used for model training at the initial time point of the prescribed image augmentation training: `noise_std:cos:0.05 → 0.015, sd_affine:cos:0.05 → 0.01, sd_deformation:linear:12.0 → 0.6, sd_simulated_bias_field:cos:0.20 → 0.03, sd_histogram_warping:cos:0.04 → 0.008`. Values for each augmentation type decrease over the course of optimization according to the prescribed schedule.

## 3 Results

We evaluate the LAMNr framework by characterizing its ability to linearize complex anatomical manifolds across two data modalities: (1) high-dimensional tabular phenotypes using RealNVP architectures, and (2) multi-modal MRI using multiscale Glow networks. Tabular experiments allow for the efficient and computationally tractable evaluation of alignment strategies, which we can then leverage to demonstrate the utility of LAMNr flows for providing a principled, likelihood-based foundation for exploring different aspects of the proposed Glow-based, LAMNr framework.

### 3.1 Tabular LAMNr Flows

Our tabular evaluation leverages the Normative Neurological Health Embedding (NNHEmbed) (Avants et al., 2025), an optimized SiMLR-based multimodal modeling framework engineered from IDPs from UK Biobank (UKBB) (Miller et al., 2016). These IDPs, comprising T1-weighted MRI (T1-w), diffusion tensor imaging (DTI), and resting-stage fMRI (rsfMRI), were generated using ANTsPyMM^5^, an ANTsX-based utility for generating tabular IDP data from neuroimaging cohorts. For NNHEmbed, the resulting views comprised 51 T1-w IDPs, 77 DTI IDPs, and 484 rsfMRI IDPs. The UKBB-based projection matrices map the multimodal (i.e., three-view) input features of 1) the Normative Neurological Library (NNL) (Gage et al., 2024) and 2) the Parkinson Progression Marker Initiative (PPMI) (Marek et al., 2011) cohorts into shared *k* = 31 dimensional bases, which then serve as the input for generating our LAMNr flows models.

#### 3.1.1 Architecture and Hyperparameter Selection

Prior to the multiview SiMLR and LAMNr flows comparative evaluation, we used the NNL and PPMI IDP data to determine optimal hyperparameter settings of the RealNVP-style normalizing flow architecture across the single modalities in terms of trained likelihoods, i.e., bits-per-dimension (BPD). The network capacity is controlled by two primary hyperparameters: (i) the coupling depth *K* (number of transform layers), and (ii) the conditioner width or the number of hidden channels (*HC*). Rather than performing an exhaustive grid search over a broad parameter space, we restricted our evaluation to a targeted window (*K* ∈ {3, 4, 5}, *HC* ∈ {64, 80, 96}). This focused selection is informed by a caution against overfitting, given the broad range in cohort sizes. An over-parameterized model risks capturing idiosyncratic noise rather than the underlying manifold geometry. By selecting the smallest architecture capable of minimizing the validation negative log-likelihood, we ensure that the model remains a compact description of the distribution.

Our hyperparameter sweep across the individual NNL and PPMI cohort views confirmed the robustness of this architectural window. For the NNL cohort (*N* = 346), a depth of *K* = 4 was consistently optimal across individual T1, DTI, and rsfMRI modalities, effectively minimizing the validation (in terms of model training) negative log-likelihood. In the PPMI cohort (*N* = 1769), while increasing capacity to *K* = 5 yielded slightly lower likelihoods, the improvement was negligible (ΔBPD < 0.001 for DTI) and did not justify the additional model complexity. Prioritizing model parsimony and methodological consistency between datasets, we fixed the configuration at *K* = 4 and *HC* = 80 for all subsequent multiview experiments. This stable parametric baseline ensures that the learned latent representations are comparable across different clinical populations while avoiding overfitting to dataset-specific noise.

#### 3.1.2 Multiview Comparison with the SiMLR NNHEmbed Framework

Building upon the baseline of individual views, we conducted a parallel evaluation of three distinct classes of latent-alignment constraints to characterize the likelihood-alignment trade-off across different mathematical frameworks. Rather than relying on a single objective, we systematically compared covariance-based (VICReg), kernel-based (HSIC), and contrastive (InfoNCE) approaches to determine how varying geometric and statistical pressures influence the resulting multiview representation. Simultaneously, we performed a systematic sweep of the alignment weight (*λ*) for each method. This allowed us to quantify the ‘likelihood penalty’ (i.e., the marginal decrease in exact BPD) as a universal cost of prioritizing a unified, multiview latent representation over view-specific details. Other related methods, such as Barlow Twins or Pearson-based correlations, were omitted as they share underlying functional principles with VICReg (specifically redundancy reduction).

Empirical results across both cohorts confirm that while all three strategies successfully align the latent manifolds, they exhibit distinct behaviors regarding density estimation and stability. HSIC emerged as the most statistically robust objective for tabular data, providing a high-fidelity alignment that effectively captures non-linear dependencies, albeit with a more pronounced likelihood penalty in the smaller NNL cohort (*BPD* = −2.871 vs. baseline −4.239). In contrast, VICReg and InfoNCE maintained likelihoods closer to the baseline (*BPD* ≈ −4.16 and −4.00 respectively for NNL), suggesting a more conservative unfolding of the anatomical manifold. Notably, in the larger PPMI dataset, the performance gap between methods narrowed significantly, with VICReg yielding a BPD (−8.142) remarkably close to the top-performaing HSIC(−8.198). This consistency, combined with its superior computational scalability, reinforces the selection of covariance-based regularization (VICReg) as the primary alignment objective for the high-dimensional imaging experiments using Glow-based LAMNr flows.

The robust latent alignment provides a biologically principled foundation for statistical inference. This refined latent space directly translates into enhanced sensitivity for clinical markers. As shown in Figure 4, LAMNr flows demonstrate significant performance gains when predicting complex cognitive and functional phenotypes compared to linear subspace projections. In the NNL cohort, the nonlinear mapping provides a substantial “correlation uplift” (Δ*r*) relative to SiMLR across multiple domains, notably in *Recall Delayed* (Δ*r* = 0.190, *q* < 10^−3^) and *Working Memory* (Δ*r* = 0.177, *q* = 0.024). In contrast, for *Reading Ability*, the nonlinear model shows a slight but non-significant decrease compared to the linear baseline (Δ*r* = −0.004, *q* = 0.960). However, when compared to an unconstrained multiview model (*λ* = 0), LAMNr alignment still retains a positive trend (Δ*r* = 0.069), suggesting that while linearity suffices for reading tasks, latent alignment remains beneficial for overall model stability. Interestingly, the performance profiles differ across populations. While the NNL cohort exhibits clear benefits from nonlinear alignment, the linear SiMLR models remain highly competitive in the PPMI cohort. This divergence likely reflects the different variance structures of the two datasets. Specifically, the NNL cohort captures a broad spectrum of healthy variation where subtle nonlinear couplings are prevalent, whereas the PPMI cohort is dominated by the strong, relatively linear pathological signal of Parkinson’s disease progression.

**Figure 4:**
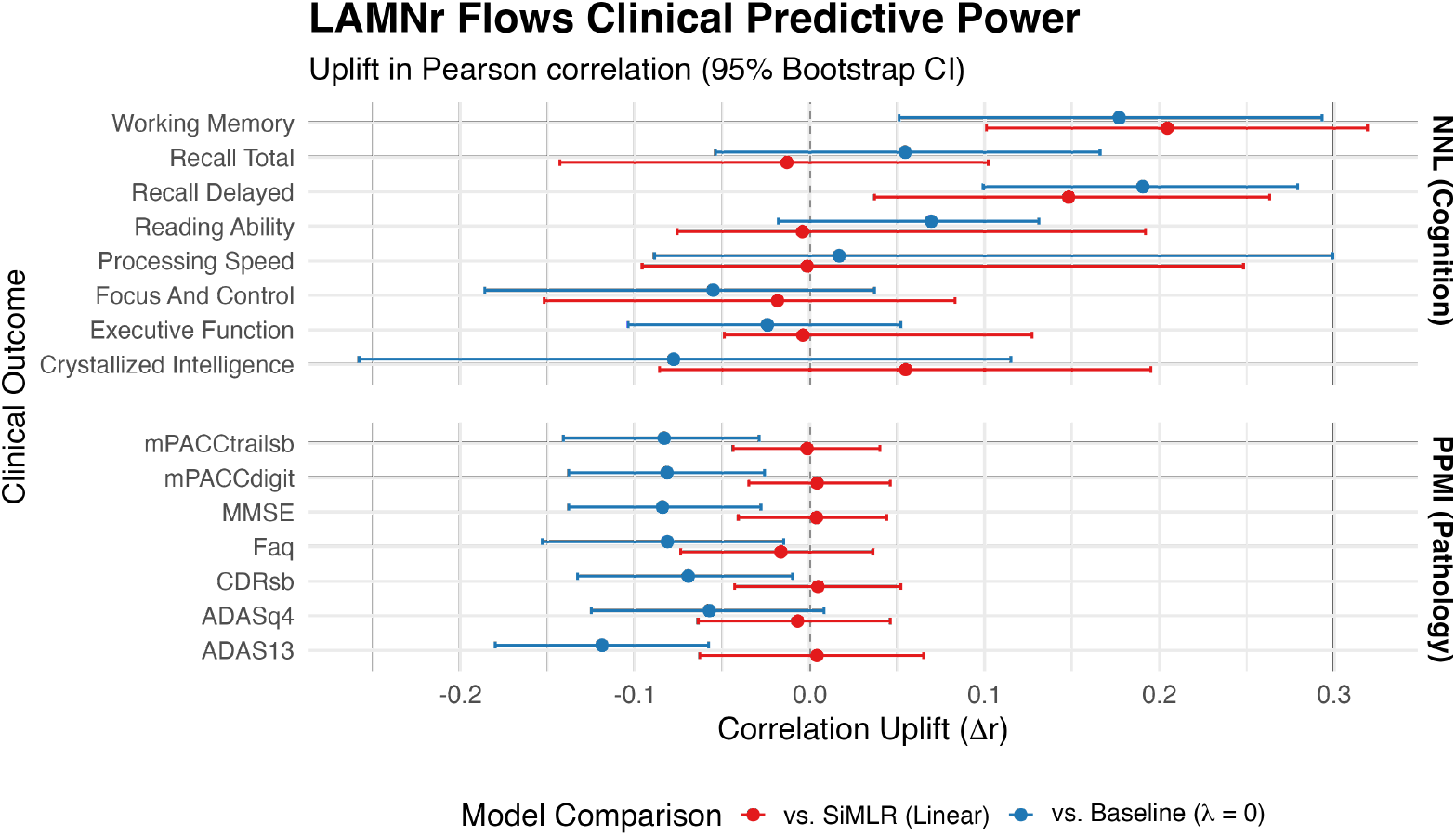
The forest plot illustrates the correlation uplift (Δ*r*) across two levels of comparison: (1) the gain from non-linear manifold mapping, represented by the difference between LAMNr flows (with VICReg) and the SiMLR baseline (i.e., red intervals), and (2) the gain from latent alignment, represented by the difference between the aligned LAMNr flows model and an unconstrained multiview baseline (*λ* = 0, i.e., blue intervals). Error bars represent the 95% confidence intervals derived from 1000 bootstrap resamples. Top panel displays results for the NNL cohort, showing significant non-linear gains in memory and executive function. Bottom panel displays results for the PPMI cohort, where linear models remain highly competitive. Significant improvements (*q* < 0.05, FDR corrected) are indicated by intervals that do not cross the zero-reference line.

### 3.2 Glow-based LAMNr Flows

#### 3.2.1 Image Data and Preprocessing

Transitioning from tabular to image data within the LAMNr Flows context introduces significant computational challenges. The Glow architecture requires storing all intermediate activations to compute exact gradients (Kingma and Dhariwal, 2018) which quickly saturates the VRAM (Video RAM), or memory, of the graphics card. This memory bottleneck is particularly acute in 3D. For instance, while a 2D slice at a standard resolution of 256^2^ contains 65,536 pixels, a corresponding 3D volume exceeds 16 million voxels. On high-capacity hardware (e.g., the NVIDIA RTX A6000 48 GB utilized here), processing volumes even at reduced resolutions like 48^3^ or 64^3^ necessitates strict architectural compromises, including reduced hidden channels and micro-batch sizes, to prevent memory overflow. Consequently, we adopt a dual organizational approach which utilizes high-resolution 2D slices as a practical sandbox for visualizing the geometric properties of the framework, while demonstrating that 3D LAMNr flows remain robust and biologically informative even at lower resolutions and structural applications. These constraints are primarily a function of current hardware availability, as the software framework is currently designed to scale with future increase in computational capabilities.

We use five data cohorts in the experiments below: the Dallas Life Brain Study (D. Park et al., 2025; D. C. Park et al., 2025), the NIMH Healthy Research Volunteer Dataset (Nugent et al., 2025), the Queensland Twin IMaging dataset (Strike et al., 2022; Strike et al., 2019), the Brain Tumor Sequence Registration Challenge dataset (Baheti et al., 2024), and the Open Access Series of Imaging Studies 3 (OASIS-3) cohort (LaMontagne et al., 2019). Whereas the first three datasets are openly available at OpenNeuro (to facilitate reproducibility), the BraTS-Reg dataset is available upon request from the challenge organizers, and OASIS-3 is available upon request through the OASIS project. These datasets are further summarized as follows:

- **Dallas Life Brain Study (DLBS)**. T1-weighted, FLAIR, diffusion-weighted MRI. Three longitudinal “waves” are included (*N*_1_ = 463, *N*_2_ = 298, *N*_3_ = 191).
- **NIMH Research Volunteer Dataset (NIMH)**. T1-weighted, T2-weighted, diffusion-weighted MRI for *N* = 234 complete subjects.
- **Queensland Twin IMaging (QTIM)**. T1-weighted MRI (*N* = 1202) including family identifiers.
- **Brain Tumor Sequence Registration Challenge (BraTS-Reg)**. T1-weighted, T1-weighted contrast enhanced, T2-weighted, FLAIR from *N* = 140 subjects, featuring preoperative and follow-up scans with expert-validated landmarks.
- **Open Access Series of Imaging Studies 3 (OASIS-3)**. Longitudinal multimodal neuroimaging (including T1-weighted MRI), clinical, and cognitive data (e.g., MMSE scores), utilizing standard FreeSurfer tabulated outputs to evaluate structural trajectories.

Common preprocessing steps for all data include rigid normalization to a common reference space, specifically the Nathan Kline Institute (NKI) template (Tustison et al., 2014) (1 mm^3^, 192 × 256 × 224) using brain-extracted T1-weighted images (Tustison et al., 2021b) and ANTs registration (Avants et al., 2014). Cropped volumetric left and right MTL sections for modeling were derived from the NIMH and OASIS-3 images using DeepFlash (Tustison et al., 2024), a deep-learning approach to parcellating specific structures of the medial temporal lobe (MTL). Similar cropped T1-w volumes from the DLBS cohort were also generated for inference. Left and right MTLs for each subject were rigidly registered independently using a label-based normalization approach (Roston et al., 2025) to the DeepFlash template (Tustison et al., 2024), oriented such that the long-axis of the hippocampus is perpendicular to the coronal plane. Fractional anisotropy (FA) images were derived from diffusion-weighted imaging using Dipy (Garyfallidis et al., 2014).

#### 3.2.2 Trained Models

To demonstrate the versatility of the LAMNr flows framework across different dimensionalities and anatomical scales, we trained three primary model configurations. These models serve as the basis for the qualitative and quantitative evaluations presented in the subsequent sections:

- **2D Multiview Model (T1-w, FLAIR, FA)**. This model was trained on mid-axial slices (index 115) from the DLBS wave 1 cohort. The architecture utilizes a 96 × 128 spatial resolution. Key hyperparameters include a multiscale depth of *L* = 5 levels, *K* = 12 coupling steps per level, and a hidden channel dimensionality of *HC* = 256. Latent alignment was enforced using VICReg (*λ* = 0.005) coupled with a CCA-based screening procedure (screening fraction = 0.5) to isolate shared anatomical features. Samples from the three modalities over the course of optimization is provided in Figure 5.
- **3D Single-view Model (T1-w)**. This whole-head model was trained on DLBS wave 1 T1-weighted volumes downsampled to 48 × 64 × 56 voxels. The architecture utilizes *L* = 3 levels, *K* = [16, 32, 64] steps, and *HC* = 96. As this represents a single-view baseline, the alignment weight was set to *λ* = 0.0, focusing purely on exact likelihood-based density estimation.
- **3D Multiview Model (T1-w, T2-w)**. This volumetric model of the left hippocampus used NIMH dataset inputs cropped to a size of 40 × 40 × 64 voxels. The network configuration consists of *L* = 3 levels with *K* = 32 coupling steps and *HC* = 128 hidden channels. Due to the high structural correlation between T1 and T2 modalities in the hippocampus, a stronger alignment weight was applied (VICReg *λ* = 1.0) with CCA-based screening.

**Figure 5:**
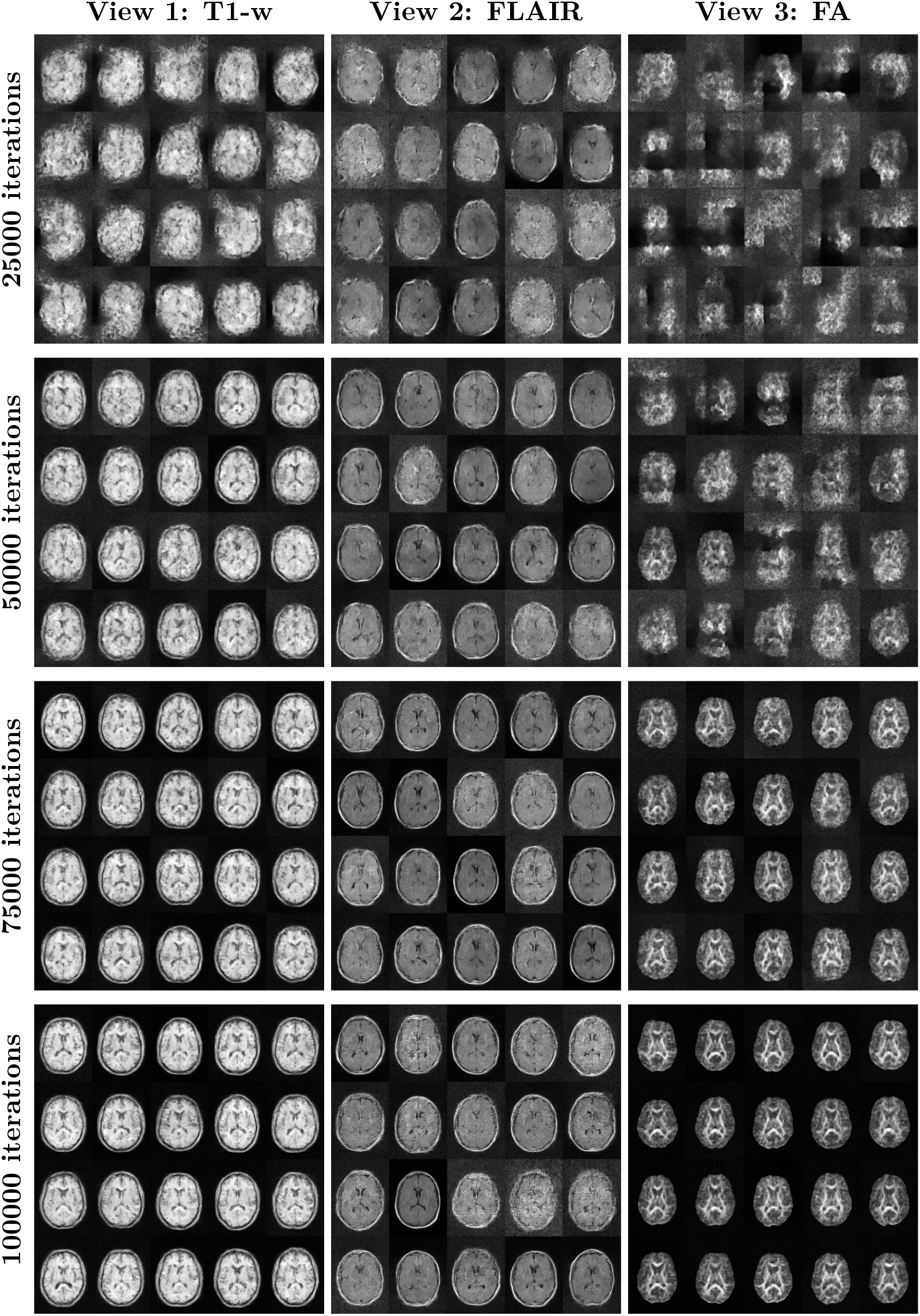
Generative samples drawn from the learned LAMNr flows model for the 3-view model (T1, FLAIR, FA) over the course of optimization. Samples were drawn at every 1000 iterations to monitor the current training state of the model. Here we show samples at 25000, 50000, 75000 and 100000 iterations. The sample temperature was *τ* = 0.8 (i.e., samples were drawn from 𝒩 (0, *τ* ^2^*I*).)

All models were optimized using the Adamax optimizer with a scheduled learning rate ranging from 2.5 × 10^−5^ to 5 × 10^−5^. To ensure numerical stability during the training of deep multiscale flows, we utilized gradient norm clipping at 0.1 or 0.2 and employed mixed precision training via a gradient scaler. To further stabilize the learned manifold and improve generative quality, an Exponential Moving Average (EMA) of model parameters was maintained throughout the optimization process. Command line interfaces for these workflows are provided through the Python-based CLIs `train_lamnr_glow_2d.py` and `train_lamnr_glow_3d.py`, which natively handle dataset-level normalization, coordinated data augmentation, and periodic logging of reconstructions for visual quality assessment.

To support downstream inference and analysis, the framework includes the `lamnr_glow_tool_2d.py` and `lamnr_glow_tool_3d.py` CLI tools, which provide extended functionality for sampling, reconstruction, and latent space manipulation. These utilities allow for the fitting of conditional Gaussian models to the learned latents via the `gauss-fit` sub-function, utilizing low-rank (SVD) or diagonal covariance estimators to bypass memory errors when processing high-dimensional image data. The `gauss-impute` sub-function facilitates the synthesis of missing modalities (e.g., T1 to FA) by leveraging the “Push-Through” Woodbury identity to ensure numerical stability in high dimensions. Furthermore, the toolkit supports the construction of population templates `(recon-template)`, latent temperature modulation to suppress anatomical anomalies `(recon-temperature)`, and geodesic interpolation that respects the high-probability manifold `(recon-interpolate)`. Finally, distances relative to the cohort mean is provided through the calculation of Mahalanobis or Euclidean distances `(calc-distance)`.

#### 3.2.3 Visualizing LAMNr Flows-based Deep Computational Anatomy

Using the 3-view model (2D) trained on the DLBS data, we demonstrate multiple applications of the LAMNr flows framework within the context of Deep Computational Anatomy (DCA). LAMNr flows provide a probabilistic, likelihood-based alternative to traditional diffeomorphometry (e.g., LDDMM), allowing for direct synthesis and manipulation of anatomical manifolds.

##### LAMNr Flows-based population template

In the context of the learned data manifold, the Fréchet mean of the anatomical distribution is efficiently approximated by decoding the origin (*z* = 0) of the latent space. By passing the zero-vector of the isotropic Gaussian prior through the inverse flow, 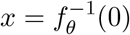, we synthesize a canonical representation that captures the central morphometric tendency of the cohort without the computational overhead of iterative diffeomorphic averaging (Avants et al., 2010). See Figure 6. Also, see `lamnr_glow_tool_2/3d.py recon-template` for more details.

**Figure 6:**
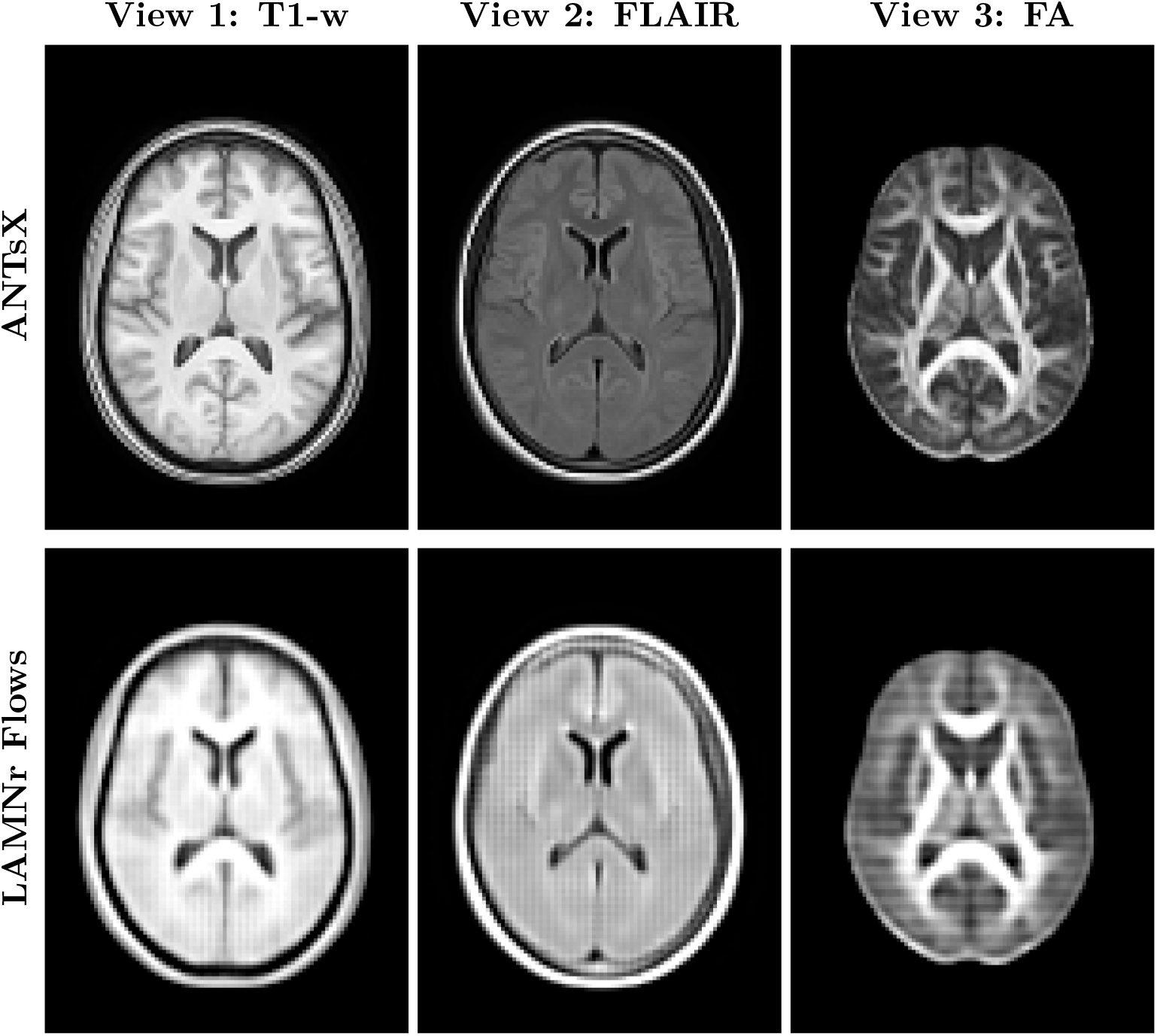
Comparison of population Fréchet mean approximations. (Top) The standard multimodal ANTsX template, constructed via traditional iterative diffeomorphic registration, representing a geometric spatial average that preserves high-frequency structural details. (Bottom) The generative latent-means, 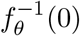, obtained in a single forward pass. The visually smoother appearance of the flow-generated template is a direct consequence of high-dimensional probabilistic modeling. As the exact mode of the latent distribution, it averages out idiosyncratic, high-frequency anatomical variations (such as specific cortical folding patterns) that do not strictly persist across the cohort. Instead of producing a single typical sample from the typical set, it models the macroscopic central morphological tendency and shared structural signal of the dataset.

##### Generative sampling

Sampling in LAMNr flows is performed by drawing a latent vector *z* from the isotropic Gaussian base distribution, *z* ~ 𝒩 (0, *τ* ^2^*I*), and mapping it back to the image domain via the inverse flow, 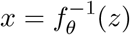. In high-dimensional latent spaces, however, the probability mass concentrates within a thin “typical set” located on a spherical shell of radius 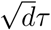, rather than near the mode at the origin. Adjusting the temperature parameter *τ* allows for explicit control over this sampling radius. Lower temperatures (*τ* < 1) contract the sampling toward the high-density (but low-volume) region near the mean to generate high-fidelity, canonical anatomies, while *τ* ≈ 1 ensures that samples are drawn from the typical set, capturing the diverse structural variations characteristic of the true empirical distribution. See Figure 7. Also, see `lamnr_glow_tool_2/3d.py` sample for more details.

**Figure 7:**
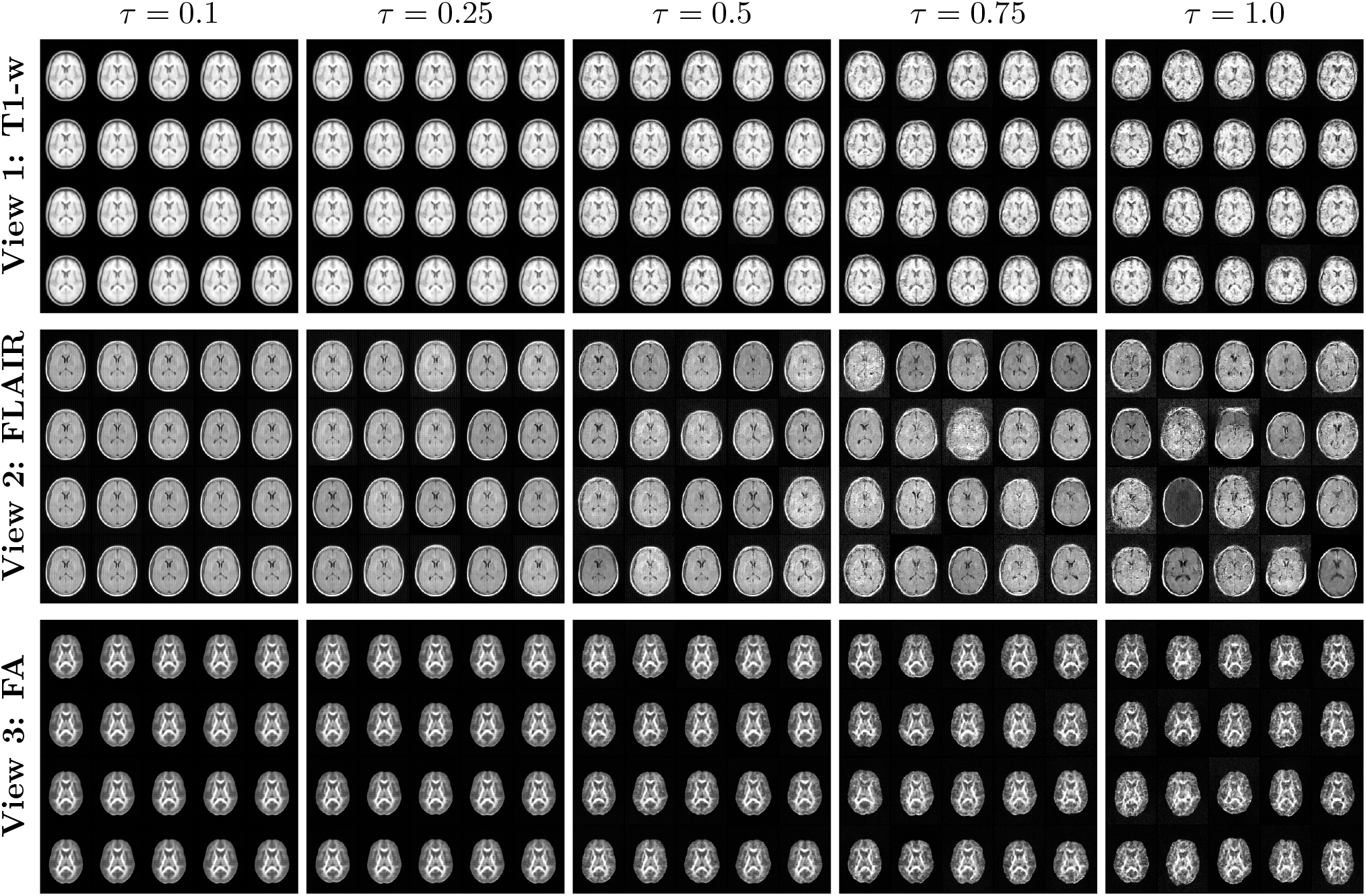
Generative samples drawn from the learned LAMNr flow prior for each view at varying temperatures (*τ*). Lower temperatures (*τ* ≤ 0.50) generate coherent and anatomically representations whereas at higher temperatures (e.g., *τ* = 1.0), the model samples extreme latent vectors from the prior tails.

##### Pairwise image interpolation

Smooth morphological transitions between anatomical scans are generated by interpolating their representations in the learned latent space. To prevent the variance collapse and out-of-distribution artifacts characteristic of standard linear interpolation, we employ a *µ*-centered spherical linear interpolation (Slerp). By applying the spherical rotation relative to the population’s empirical mean, *µ*, the latent trajectory better preserves the intrinsic data variance, ensuring all intermediate representations remain closer to the high-probability anatomical manifold.

This is demonstrated in Figure 8 for both within-cohort data (i.e., DLBS Wave 2) and out-of-cohort data (i.e., BraTS-Reg), in terms of the model training data (i.e., DLBS Wave 1). We observe smooth transitions for both the T1-w and FLAIR images between the source (*t* = 0.0) and target images (*t* = 1.0). In the case of the DLBS Wave 2 cohort, we selected two subjects of different ages which illustrates the morphological interpolation from larger to smaller ventricles and from the presence to absence of white matter hyperintensities. We see similar high quality interpolations in a BraTS-Reg subject (Subject 5, post- and pre-resection scans). It is noteworthy reiterating that training data did not include skull-stripped images. See `lamnr_glow_tool_2/3d.py recon-interpolate` for specific implementation details.

**Figure 8:**
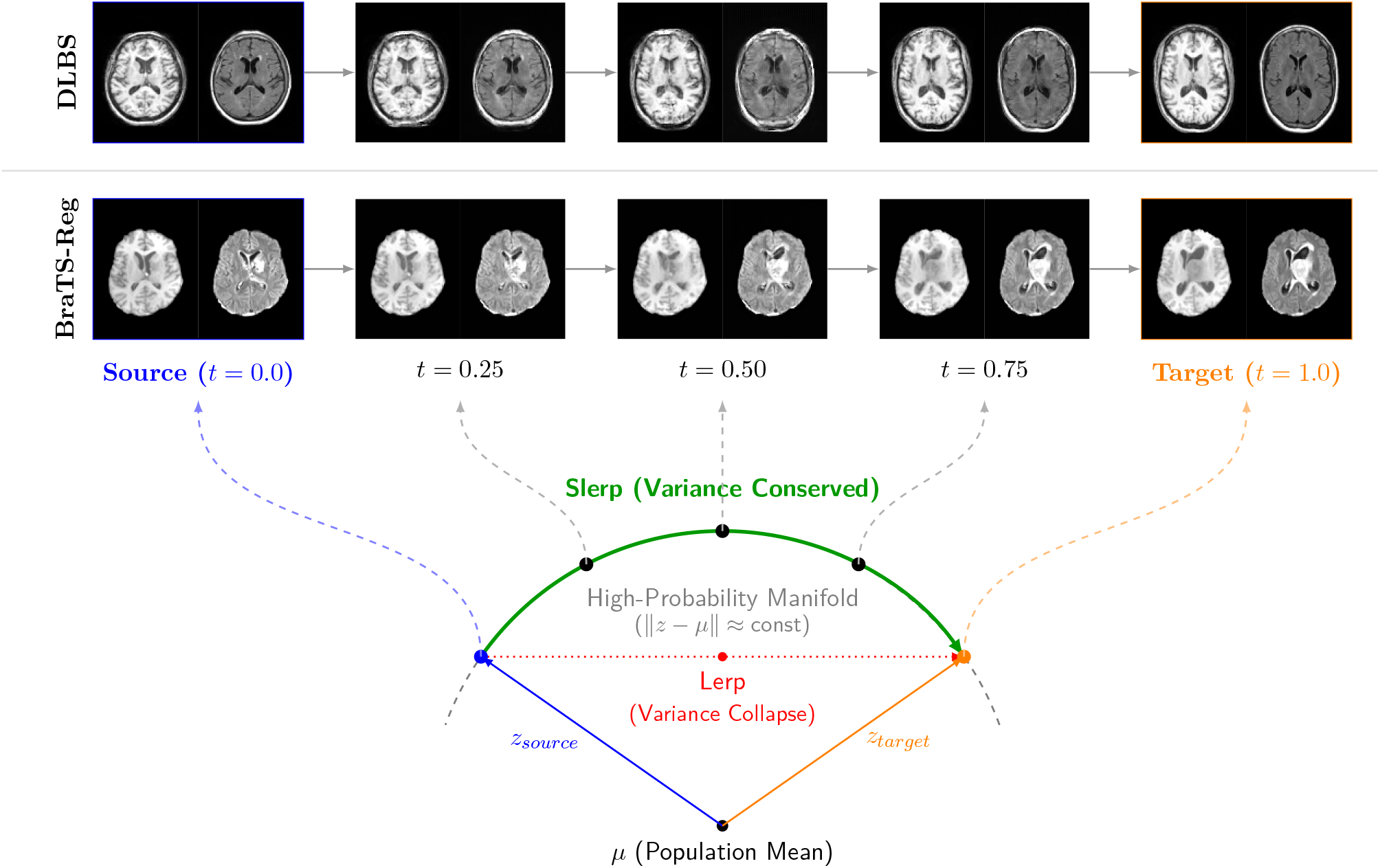
Interpolation using the DLBS Wave 2 cohort (top row) and BraTS-Reg cohort (second row). Model training used only whole-head DLBS Wave 1 data (T1-w, FLAIR, FA). (Top) The generated morphological transition between a source image (*t* = 1.0) and a target (*t* = 0.0) multimodal images (T1-w, FLAIR). Interpolation DLBS data (Wave 2) included the source image (Subject 4488, Age 77) and target image (Subject 587, Age 53). BraTS-Reg is demonstrated using pre- and post-resection T1-w and FLAIR images from Subject 5. (Bottom) A geometric representation of the joint latent space. The empirical distribution of the training cohort is centered around *µ*. Standard linear interpolation (Lerp, dotted red line) cuts through the interior of the latent hypersphere, causing a severe contraction of the vector’s norm (variance collapse). This forces the decoding flow to evaluate out-of-distribution coordinates. Conversely, applying Slerp relative to the empirical mean *µ* (solid green arc) better preserves the natural variance of the data such that the trajectory follows the high-probability manifold.

##### Cross-modal imputation via Conditional Gaussian modeling

Missing modalities are synthesized by encoding the available observed images to the latent space, *z*_*O*_ = *f* ^(*O*)^(𝒩 ^(*O*)^), and computing the exact conditional expectation of the unobserved latent vectors, *µ*_*U*|*O*_, under the learned joint Gaussian prior. Projecting this optimal estimate through the target modalities’ inverse flows, 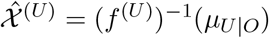, yields a high-fidelity imputation that guarantees mathematical consistency with the population’s cross-view dependencies. Crucially, because the joint prior models the full multi-view latent space simultaneously, this formulation is inherently flexible. It supports conditioning on any arbitrary subset of available data, enabling complex many-to-many translations (e.g., synthesizing a single FA map from combined T1-w and T2-w inputs, or simultaneously generating T2-w and FA from a single T1-w scan). See Figure 9. Also, see `lamnr_glow_tool_2/3d.py impute` for more details.

**Figure 9:**
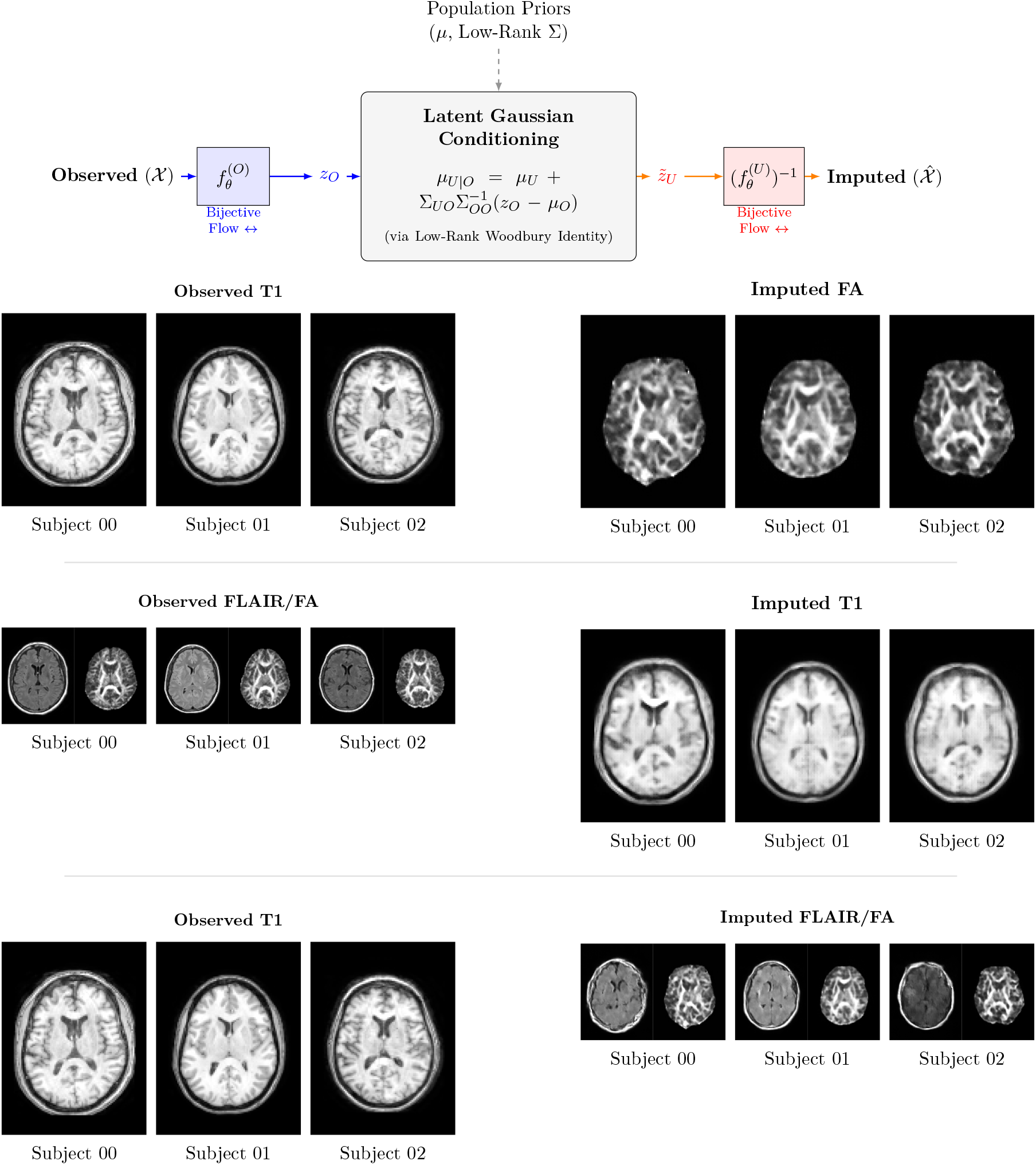
(Top) Diagrammatic illustration of the Conditional Gaussian modeling approach available through the LAMNr flows framework. Observed input features 𝒳 are mapped to the latent representation *z*_*O*_ through the learned bijective flow *f*_*θ*_. Imputation of missing modalities is performed via latent Gaussian conditioning modeling. The target image 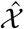 is synthesized by projecting the imputed latent vector 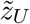 back to the data space via the inverse flow 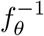. (Bottom) Performance is demonstrated across three subjects under varying observational constraints. (Row 1) Synthesis of Fractional Anisotropy (FA) maps from observed T1-weighted inputs. (Row 2) Joint reconstruction of T1-weighted scans from observed FLAIR and FA modalities. (Row 3) Simultaneous multi-modal imputation of FLAIR and FA from a single observed T1 input.

##### Latent distances

The bijective nature of normalizing flows allows complex anatomical deviations to be quantified through a flexible suite of distance metrics in the learned latent space, depending on the analytical objective. Euclidean distance provides a straightforward measure of separation for basic similarity assessments. To account for the natural variance of each latent dimension, we implement a standardized Euclidean (diagonal Mahalanobis) distance, 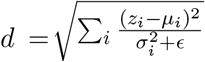, which benchmarks a subject against the normative Gaussian mean (*µ*) without artificially penalizing high-variance anatomical traits. For point-to-point comparisons between the latents *z*_*j*_ and *z*_*k*_ of specific images, we utilize geodesic distance derived from cosine similarity, *d* = arccos(clamp(sim(*z*_*j*_, *z*_*k*_))). By measuring the angular displacement on the hypersphere, this metric respects the spherical geometry of the isotropic Gaussian prior, ensuring that anatomical transitions are evaluated along the high-density manifold. These combined metrics yield a rigorous, variance-weighted framework for anomaly detection and longitudinal assessment. See Figure 10. Also, see `lamnr_glow_tool_2/3d.py calc-distance` for more details.

**Figure 10:**
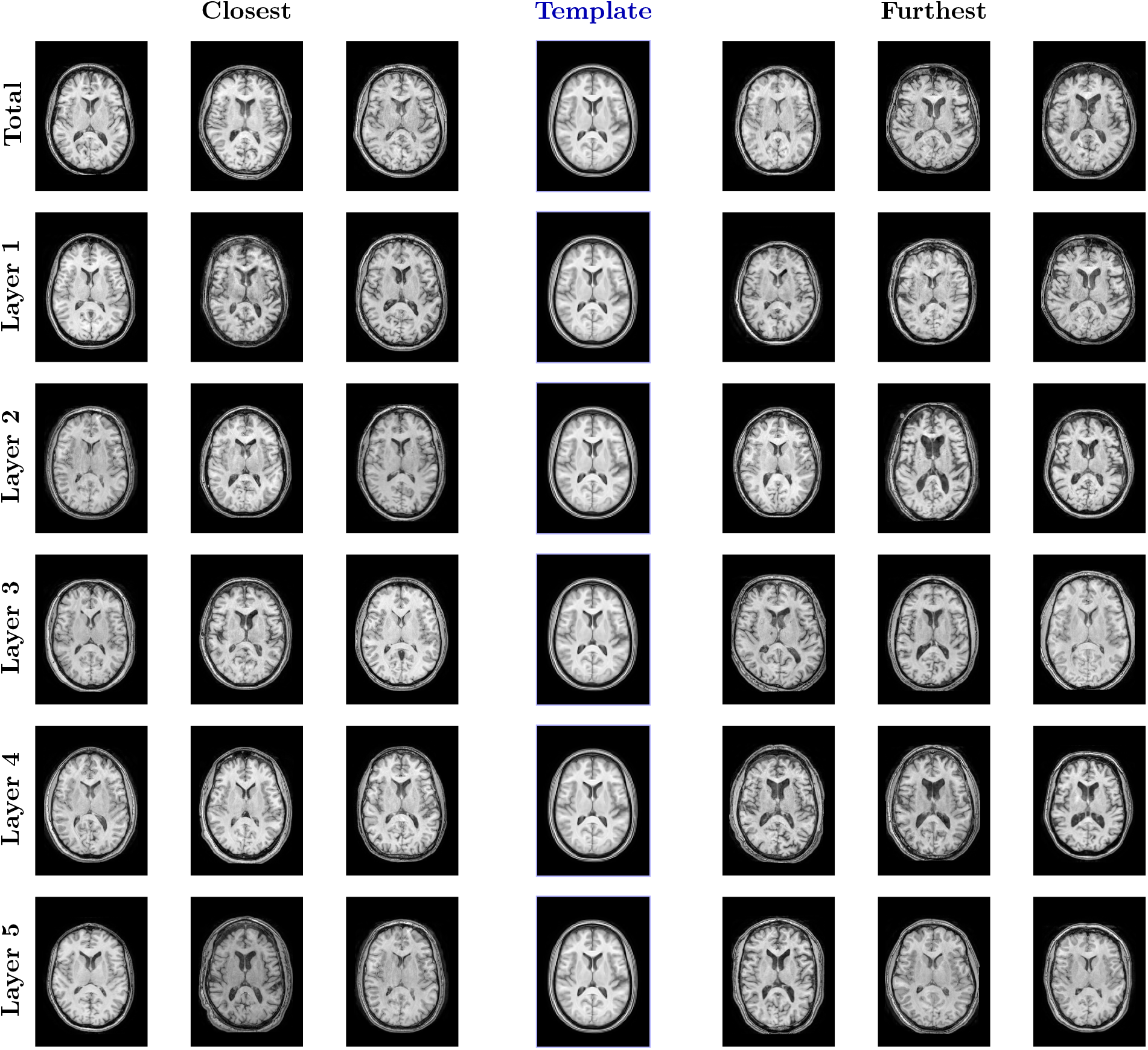
Visualization of latent space distance (cf Equation 8) across the DLBS Wave 2 cohort with respect to the ANTsX T1-w template (cf Figure 6). We calculate the total latent space distance and the distance for each hierarchical Glow layer and render the closest images (left) and the furthest images (right) centralizing the template as a visual reference point.

**Figure 11:**
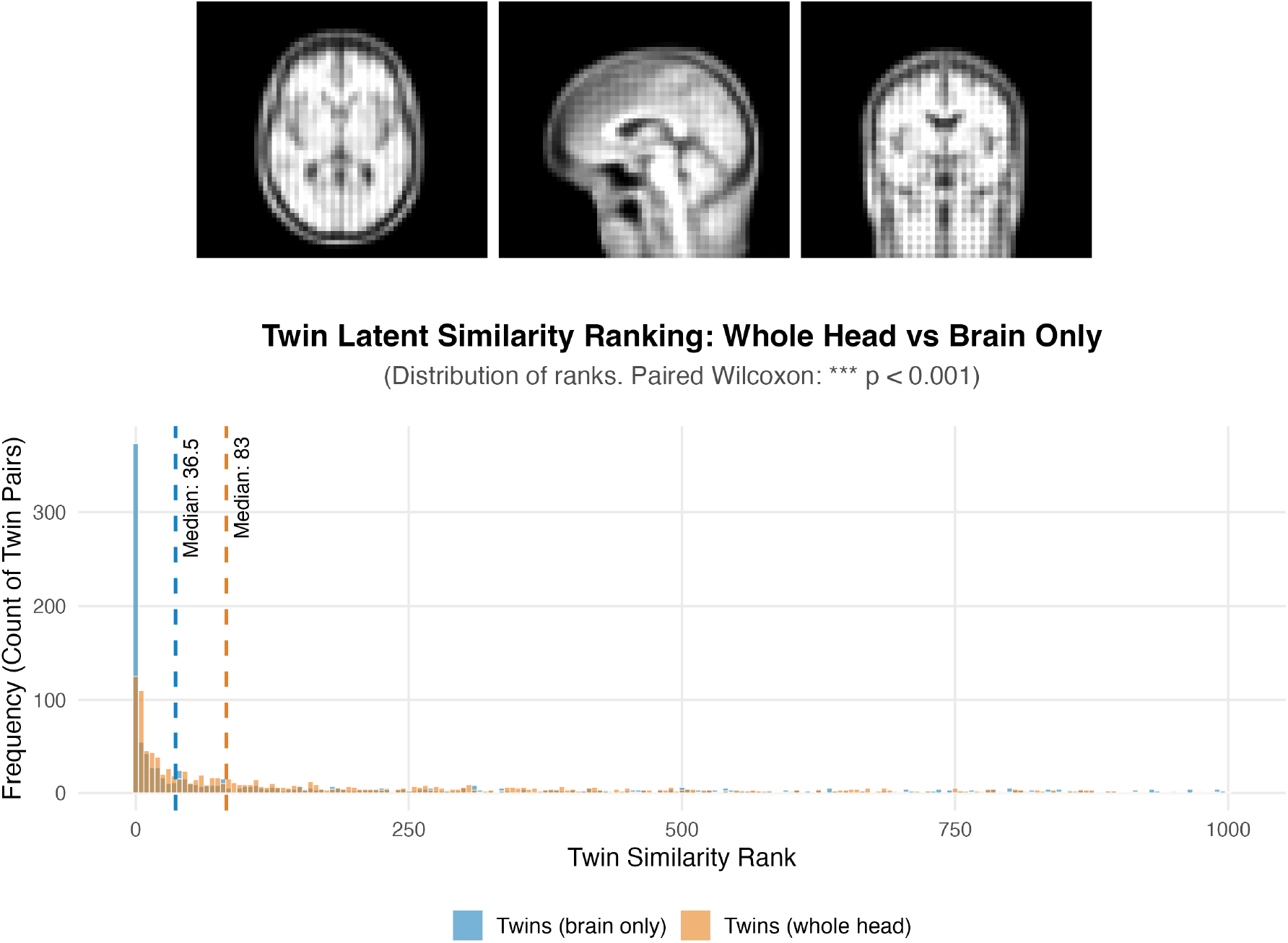
Top row: Canonical views of the latent-defined template from the 3D, T1-w volumetric LAMNr flow model constructed from the DLBS wave 1 data. Second row: Distribution of latent similarity ranks between twins (with and without brain extraction) from the QTIM dataset. The latent distance was calculated from each subject to every other subject for which a ranking was derived per subject. A rank closer to 1 indicates the highest possible similarity (i.e., indicative of the twin counterpart). The overlaid distributions compare imaging data including the effects of brain extraction. The vertical dashed lines indicate the respective medians of the two groups. Skull-stripped images significantly lower the median similarity rank to 36.5, compared to a median rank of 83 for whole-head images (*p* < 0.001).

#### 3.2.4 T1-w Volumetric LAMNr Flows Model

The utility of low resolution 3D single-view, T1-w LAMNr flows models was demonstrated on the QTIM (Twins) dataset. It was hypothesized that the latent distance could serve as a similarity index for predicting twin pairs. Although it was trained on whole-head data (DLBS, wave 1), we also demonstrate that it generalizes to skull-stripped data (cf. Figure 8) and, in fact, this preprocessing step improves prediction performance. T1-weighted imaging volumes from the QTIM cohort were analyzed under two preprocessing conditions: original full-head volumes and skull-stripped volumes isolating the brain parenchyma (Tustison et al., 2021b). Images were projected into the LAMNr flows latent space, and an *N* × *N* inter-subject distance matrix was generated to evaluate anatomical affinity by converting these distances into similarity ranks. A rank of 1 indicates that a subject’s twin is their nearest neighbor in the latent space. Statistical results demonstrate significantly higher discriminative power when the model processes brain-only images, with the median similarity rank for twin pairs improving from 83 (whole head) to 36.5 (brain only). A paired Wilcoxon test confirmed that this improvement is significant (*p* < 0.001).

#### 3.2.5 Multiview Modeling of the Medial Temporal Lobe

We evaluated the ability of the proposed framework to capture clinically relevant changes of the medial temporal lobe (MTL). The LAMNr flows model is volumetric (40×40×64 voxels) comprising two views/modalities (T1-w, T2-w) common to imaging of the MTL (Wisse et al., 2017; Yushkevich et al., 2015). T1-w/T2-w imaging pairs (*N* = 249) were taken from the NIMH dataset (Nugent et al., 2025). Training included both the left and right (flipped) MTLs. The cropped volume for each image set was defined by applying DeepFLASH (Tustison et al., 2024), an MTL segmentation application available in ANTsXNet and ANTsTorch, to linearly normalize each hemisphere to the DeepFLASH template with the long axis of the hippocampus oriented perpendicular to the coronal plane (see Figure 12).

**Figure 12:**
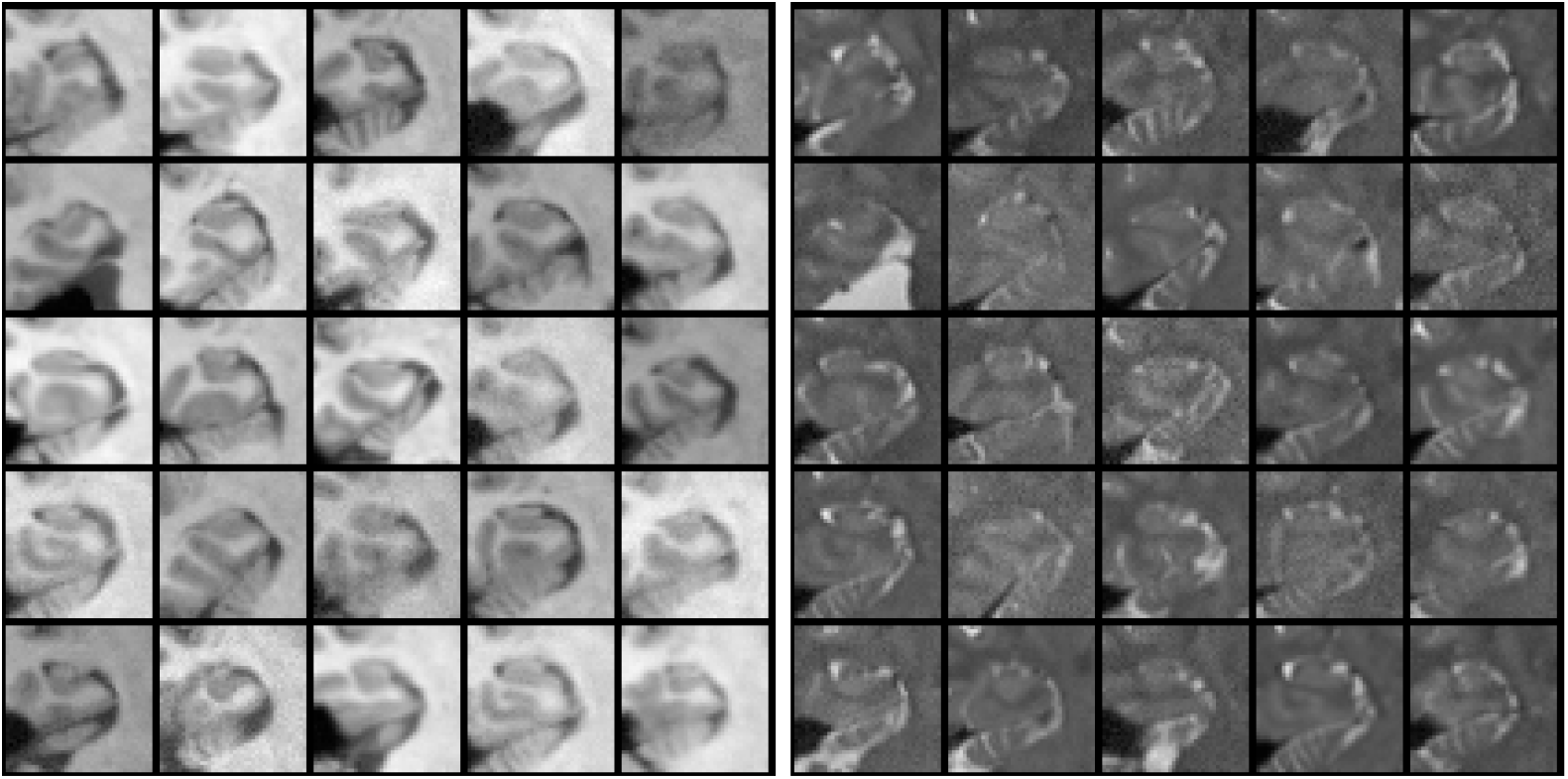
Representative 2D oblique slices from the 3D T1-weighted (left) and T2-weighted (right) medial temporal lobe (MTL) training pairs from the NIMH dataset. The volumes have been cropped and rigidly normalized to the DeepFLASH template, orienting the longitudinal axis of the hippocampus perpendicular to the coronal plane.

Each view-specific Glow instance was scaled to maximize spatial expressivity while adhering to hardware constraints, utilizing three multiscale resolution levels (*L* = 3), 32 coupling steps per level (*K* = 32), and 128 hidden channels. Training was stabilized using an effective batch size of 64 (`BATCH=8, GRAD_ACCUM=8`). The data augmentation schedule was: `noise_std: cos:0.05->0.02, sd_deformation:linear: 6.0->0.2, sd_simulated_bias_field: cos:0.20->0.01`, and sd_histogram_warping: cos:0.04->0.002 over 80000 iterations with a total of 100000 iterations. To ensure anatomical synchronization between modalities without overriding their respective characteristics, multimodal regularization was applied to the latent space. Alignment was jointly driven by VICReg to maintain the invariance of shared structural representations between T1-w and T2-w views, while penalizing variance to prevent dimensional collapse of the latent space. Specifically, the VICReg objective was configured with an overall alignment weight of 1.0, utilizing penalty coefficients of 25.0 for invariance, 25.0 for variance, and 1.0 for covariance, alongside a variance hinge threshold (*γ*) of 1.0. Additionally, Canonical Correlation Analysis (CCA) screening was dynamically deployed to isolate and project highly correlated cross-modality features. The CCA screening was activated after a warmup period of 1000 iterations and refreshed every 5000 iterations. To ensure robust feature selection, the screening retained the top 50% of the features (SCREEN_FRAC=0.5, PREFILTER_FRAC=0.5) and applied a ridge penalty of 10^−3^ to stabilize the covariance matrix inversion.

To validate the clinical utility of the learned structural representations, we evaluated the LAMNr latent space using longitudinal data from the OASIS-3 cohort (LaMontagne et al., 2019) which includes standard FreeSurfer output compiled in tabular form. Intra-subject structural trajectories were quantified with our LAMNr flows approach by calculating the spherical linear interpolation (Slerp) geodesic distance in the latent space between a subject’s baseline scan and subsequent followup visits for both the left and right (flipped) MTLs. We compared the resulting statistical model of these latent geometric deformations against a FreeSurfer composite volumetric biomarker (hippocampus, entorhinal cortex, and parahippocampal cortex) for modeling cognitive decline (Schwarz et al., 2016), measured via the Mini-Mental State Examination (MMSE).

A linear mixed-effects (LME) model incorporating the standardized geodesic distance, age at visit, and intracranial volume (ICV) as fixed effects, with a random intercept for each subject, was formulated to evaluate the longitudinal trajectories. Specifically, the LAMNr spatial deformation model is defined as:

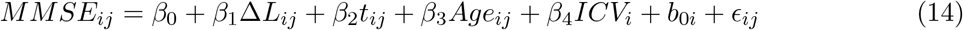

where *MMSE*_*ij*_ is the clinical cognitive score for subject *i* at visit *j*, Δ*L*_*ij*_ is the standardized Slerp geodesic distance from the subject’s baseline latent representation, *t*_*ij*_ represents the longitudinal time elapsed (in years) since the baseline scan, *Age*_*ij*_ is the standardized age at the time of the visit, and *ICV*_*i*_ is the standardized intracranial volume. The term 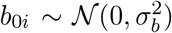 represents the subject-specific random intercept accounting for baseline cognitive variability, and *ϵ*_*ij*_ ~ 𝒩 (0, *σ*^2^) is the residual error.

Analysis of the cohort follow-up data revealed that longitudinal divergence along LAMNr geodesic trajectories is significantly, and negatively, associated with cognitive performance (*β* = −0.142, *p* < 0.01). Specifically, greater geodesic distance from a subject’s baseline representation corresponds to a steeper decline in MMSE scores. Notably, when stratifying the cohort to evaluate preclinical sensitivity (i.e., restricting the analysis exclusively to subjects clinically diagnosed as cognitively normal) the latent distance remained a robust predictor of subtle MMSE variations (*p* < 0.001). This indicates that the proposed approach potentially captures early, sub-macroscopic morphological shifts in the MTL that precede the gross volumetric tissue loss traditionally isolated by macroscopic segmentation workflows.

For methodological comparison, the macroscopic volumetric model substitutes the latent geodesic distance with the standardized FreeSurfer composite AD signature volume (*V*_*ij*_):

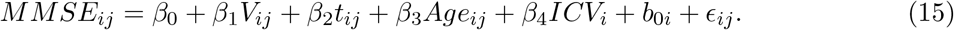

As anticipated for a targeted macroscopic biomarker, the FreeSurfer composite AD signature demonstrated a robust positive association with cognitive performance (*β* = 0.830, *p* < 0.001), confirming that gross volumetric atrophy in these regions strongly parallels clinical decline. To contextualize these findings against standard macroscopic techniques, we compared the global model fits. While the FreeSurfer composite AD formulation yielded a lower Akaike Information Criterion (AIC = 4290.3) than the LAMNr geodesic trajectory model (AIC = 4389.9) across the full follow-up cohort, this performance differential is methodologically consistent. FreeSurfer relies on explicit spatial priors and supervised atlas-based segmentations specifically engineered to isolate the macroscopic epicenters of Alzheimer’s disease pathology. Conversely, the LAMNr framework is entirely unsupervised and data-driven, lacking explicit anatomical priors.

## 4 Discussion

LAMNr flows provide a flexible framework for data modeling which provides exact likelihoods and bijective mappings between input spaces and their corresponding latent spaces. Among other possibilities, this permits a deep learning-based perspective of traditional CA. Historically, traditional CA has been fundamentally defined by image registration in which diffeomorphic transformation groups are leveraged to model biological shape variability. DCA, as described and implemented within the LAMNr flows framework, bypasses this explicit image registration requirement by topologically unfolding non-linear anatomical manifolds into a structured latent space via coordinated normalizing flows instances. Instead of iterative registration workflows, DCA employs single-pass bijective mappings per modality for inferring fundamental CA concepts.

As a salient example, traditional population template construction requires image registration for establishing joint anatomical correspondence as a prerequisite for computing the central intensity and morphological tendency of a cohort. In contrast, the DCA-based population template is defined as the inverse mapping of the latent origin which provides a barycentric anchor in latent space for characterizing the cohort central tendency. Because this generative template is the exact mode of the learned distribution, it naturally filters idiosyncratic high-frequency noise, yielding a smooth representation of shared structural signals. Furthermore, to navigate this space without the variance collapse typical of high-dimensional Euclidean operations, we utilize spherical linear interpolation. This ensures that interpolative trajectories remain strictly on the typical set, i.e., the high-probability manifold where realistic anatomical instances reside. Similarly, the metric operations of DCA substitute the image registration of CA with algebraic interpolations calculated directly within the latent manifold which respect the underlying latent-space geometry with efficient inverse single passes through the network(s). By leveraging the exact-likelihood foundations of certain normalizing flows architectures, this framework parallels the geometric rigor of traditional CA while potentially providing a deep learning-based approach for multimodal biological analysis.

Beyond CA modeling, the empirical results demonstrate the utility of LAMNr flows for clinical and biological investigation. In the tabular experiments, LAMNr flows achieved a significant “correlation uplift” over linear SiMLR baselines when predicting cognitive outcomes such as working memory and delayed recall within the NNL cohort. This suggests that the non-linear unfolding of the anatomical manifold captures subtle biological couplings that are inaccessible to linear subspace projections. However, the competitive performance of linear models in the PPMI cohort indicates that pathological signals, such as those associated with Parkinson’s disease, may be dominated by stronger, more linear variance structures. This divergence highlights the importance of selecting alignment strategies, such as VICReg or HSIC, that balance density estimation with the specific geometric attributes of the datasets of interest.

While traditional diffeomorphic image registration algorithms excel at alignment for large deformation scenarios, significant topological disruptions, such as tumor-induced changes, can limit accuracy. One of our early hypotheses in the development of this work was that DCA-based latent interpolation would be able to overcome such topological difficulties by providing an intermediate image (*t* = 0.5) for more robust image registration, in the style of classic Symmetric Normalization. The BraTS-Reg22 Challenge (Baheti et al., 2024) data provided a potentially ideal opportunity to test such an hypothesis as it involves pre- and post-resection data with expert-annotated landmarks. Although preliminary evaluations demonstrated competitive accuracy (cf. Figure 8), the limited resolution of our 3D LAMNr models was insufficient for the task and will be addressed in future work.

Our longitudinal MTL analysis demonstrates that unsupervised, multiview representations can effectively model disease progression without regional priors. The fact that LAMNr achieves highly significant longitudinal predictive power (*β* = −0.142, *p* = 0.008) based solely on latent spatial distances highlights its capacity to encode relevant neurodegenerative trajectories intrinsically. Crucially, we observe a temporal divergence in biomarker utility. While macroscopic volumetry dominates the later stages of gross tissue loss, which accounts for the superior global fit (AIC) of the FreeSurfer composite model on the full OASIS-3 cohort. By identifying structural deviations in subjects clinically diagnosed as cognitively normal, the LAMNr framework potentially captures the early, sub-macroscopic morphological shifts that precede the overt atrophy required for conventional segmentation approaches.

Finally, we must explicitly acknowledge the computational limitations of the current work. While the LAMNr flows framework provides a mathematically rigorous foundation for deep computational anatomy, the underlying Glow-style architectures are inherently memory-intensive due to the requirement of storing intermediate activations for exact gradient computation. Currently, this VRAM bottleneck restricts the application of high-resolution 3D models to targeted sub-structures (e.g., the MTL) or necessitates the downsampling of whole-brain volumes. However, the framework is currently effective for both higher-resolution 2D exploration and localized 3D applications. Furthermore, our comprehensive open-source software implementation, natively integrated into the ANTsX ecosystem via ANTsTorch, is fully 2D and 3D capable by design. To mitigate existing constraints, we have implemented architectural refinements such as gradient microbatching, bounded coupling scales, and a low-rank-plus-diagonal Gaussian parameterization (solved via the Woodbury matrix identity) to enable exact conditional inference for cross-modal imputation in 3D. Our ongoing and future research is actively focused on remedying these memory limitations. By exploring advanced algorithmic efficiencies and leveraging next-generation hardware, we aim to extend this exact-likelihood framework to high-resolution, whole-brain 3D volumes, providing a robust and scalable foundation for computational anatomy.

## Acknowledgments

Support for the research reported in this work includes funding from the Office of Naval Research (N0014-23-1-2317) and the National Institute of Biomedical Imaging and Bioengineering (R01-EB031722).

## Footnotes

1 In practice, we choose the alignment objective based on both computational budget and the expected cross-view structure. For small batches or limited computational resources, simple second-order methods such as Pearson correlation, or non-contrastive objectives such as VICReg, are attractive because they are stable and inexpensive to estimate. When stronger redundancy reduction is needed while still avoiding negative pairs, Barlow Twins is a good default, explicitly driving cross-correlation matrices towards identity. InfoNCE is most useful when we can train with large batches and many in-batch negatives, for example in discriminative multiview matching or retrieval-style scenarios where each sample in one view must be matched against many candidates in another. HSIC-based alignment is reserved for settings where we expect predominantly non-linear cross-modality relations as it can capture richer dependencies than correlations but carries an *O*(*B*^2^) kernel cost and is more sensitive to kernel and bandwidth choices.

2 In some early experiments we replaced the fixed weighting, *λ*, by learned task weights following the homoscedastic aleatoric uncertainty scheme of Kendall and Gal (Kendall et al., 2018). In this formulation, each loss term *L*_*i*_ is scaled as 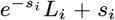, where 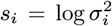 represents task-dependent aleatoric (data) uncertainty, as opposed to epistemic (model) uncertainty (Hüllermeier and Waegeman, 2021). While this can automatically balance losses with different units, in our multiview setting it tended to inflate the alignment variance and drive the effective alignment weight 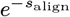 toward zero, effectively suppressing latent alignment. Therefore, we report results using a fixed *λ* schedule in the main experiments.

3 Recent transformer autoregressive flows such as STARFlow achieve strong high-resolution synthesis by operating as a normalizing flow in the latent space of a pretrained autoencoder (Gu et al., 2025). This design does not provide an exact, per-sample bijection from pixel space to the flow’s latents or multiscale per-level latents for analysis, both of which we require for per-level alignment and post-hoc Gaussian conditioning. This motivates our adoption of Glow-style multiscale flows that offer single-pass, exact encoding/decoding in image space with explicit latent access (Kingma and Dhariwal, 2018).

4 https://www.inference.vc/high-dimensional-gaussian-distributions-are-soap-bubble/

5 https://github.com/ANTsX/ANTsPyMM

